# Cell – ECM interactions play distinct and essential roles at multiple stages during the development of the aortic arch

**DOI:** 10.1101/2020.04.07.029926

**Authors:** Michael Warkala, Dongying Chen, Ali Jubran, AnnJosette Ramirez, Michael Schonning, Xia Wang, Huaning Zhao, Sophie Astrof

## Abstract

**Rationale:** Defects in the morphogenesis of the 4^th^ pharyngeal arch arteries (PAAs) give rise to lethal birth defects. Understanding genes and mechanisms regulating PAA formation will provide important insights into the etiology and treatments for congenital heart disease.

**Objective:** Cell-ECM interactions play essential roles in the morphogenesis of PAAs and their derivatives, the aortic arch artery (AAA) and its major branches; however, their specific functions are not well-understood. Previously, we demonstrated that integrin α5β1 and fibronectin (Fn1) expressed in the *Isl1* lineages regulate PAA formation. The objective of these studies was to investigate cellular mechanisms by which integrin α5β1 and Fn1 regulate AAA morphogenesis.

**Methods and Results:** Using temporal lineage tracing, whole-mount confocal imaging, and quantitative analysis of the second heart field (SHF) and endothelial cell (EC) dynamics, we show that the majority of PAA EC progenitors arise by E7.5 in the SHF and populate pharyngeal arch mesenchyme between E7.5 and E9.5. Consequently, SHF-derived ECs in the pharyngeal arches become organized into a uniform plexus of small blood vessels, which becomes remodeled into the PAAs between 31 – 35 somites. The remodeling of the vascular plexus is orchestrated by signals dependent on pharyngeal ECM microenvironment extrinsic to the endothelium. Conditional ablation of integrin α5β1 or Fn1 in the Isl1 lineages showed that signaling by the ECM regulates AAA morphogenesis at multiple steps: 1) the recruitment of the SHF-derived ECs into the pharyngeal arches, 2) the remodeling of the uniform EC plexus in the 4^th^ arches into the PAAs; and 3) differentiation of neural crest-derived cells abutting the PAA endothelium into vascular smooth muscle cells.

**Conclusions:** PAA formation is a multi-step process entailing dynamic contribution of SHF-derived ECs to pharyngeal arches, the remodeling of endothelial plexus into the PAAs, and the remodeling of the PAAs into the AAA and its major branches. Cell-ECM interactions regulated by integrin α5β1 and Fn1 play essential roles at each of these developmental stages.

## Introduction

Aortic arch artery (AAA) and its major branches comprise an asymmetrical vascular tree that routes oxygenated blood from the heart into the systemic circulation ^1^. Defects in the development of the AAA cause devastating forms of congenital heart disease (CHD) due to interruption(s) in the aortic arch, of which interrupted aortic arch type B (IAA-B) is more prevalent ^2, 3^. Non-lethal defects in aortic arch morphogenesis such as vascular rings can impact the quality of life by causing constriction of the trachea and esophagus, and resulting in difficulties with eating, breathing, and also in dizziness, vertigo, or tinnitus ^4^.

The AAA and its major branches develop from the remodeling of three bilaterally symmetrical pairs of pharyngeal arch arteries (PAA), numbered 3, 4, and 6 ^5^. It is important to note that phenotypically identical AAA defects arise due to either defects in PAA formation or defects in the remodeling of initially well-formed, symmetrical PAAs into asymmetric AAAs ^6^. PAAs arise by vasculogenesis from endothelial precursors originating in the lateral plate mesoderm, also known as the second heart field (SHF) ^7-13^. Experiments in zebrafish and mice have demonstrated that PAA formation is a multi-stage process that entails endothelial specification in the SHF, migration of SHF-derived endothelial progenitors into the pharyngeal region, differentiation into ECs, and the assembly of SHF-derived ECs into a plexus of small blood vessels ^9, 13-16^. Thereafter, the pharyngeal endothelial plexus becomes connected with the ventral and dorsal aortae. The endothelium of the ventral aortae also forms by vasculogenesis from SHF-derived progenitors and is contiguous with the cardiac outflow tract and the PAAs ^9, 11^. Following pharyngeal arch segmentation, the plexus endothelium within each arch is rearranged into the PAAs ^9^. The 3^rd^ PAA is evident by E9.5, before the 4^th^ and 6^th^ PAAs are formed. By the evening of E10.5, all three symmetrical pairs of PAAs are formed. Defects in the formation of the left 4^th^ PAA lead to IAA-B, which is lethal unless corrected by surgery soon after birth ^2^. Following PAA formation, neural crest-derived cells closest to the PAA endothelium differentiate into vascular smooth muscle cells (VSMCs), surrounding the PAA endothelium with a VSMC coat by E12.5 ^17-21^. While not essential for PAA formation, the differentiation of neural crest (NC)-derived cells into VSMCs is essential for the stability of the PAAs, and for their eventual remodeling into the asymmetrical AAA and its branches; Defects in NC differentiation in the 4^th^ pharyngeal arch lead to arch artery regression, and IAA-B ^19, 20, 22^. In summary, IAA-B results due to either defects in the formation of the left 4^th^ PAA or due to its regression.

Morphogenesis of distinct organs and structures proceeds within niches comprised of distinct complements of extracellular matrix (ECM) proteins, and alterations in ECM microenvironment can severely affect embryogenesis ^23-27^. We discovered that the pharyngeal arch microenvironment is enriched in the ECM glycoprotein fibronectin (Fn1) both at the mRNA and protein levels ^28^. Fn1 is highly expressed in the pharyngeal endoderm, ectoderm, endothelium, and the second heart field (SHF) mesoderm between E8.5 and E10.5, the period coinciding with PA formation ^28, 29^. Between E10.5 and E11.5 Fn1 becomes highly upregulated in the NC-derived cells abutting the 4^th^ PAA endothelium, corresponding with the time these cells differentiate into VSMCs. Our previous studies demonstrated that local depletion of Fn1 in the pharyngeal microenvironment using the Isl1^Cre^ knockin mice or in the NC-derived cells, using a variety of NC-expressing Cre lines, resulted in the IAA-B and RERSA ^29, 30^. However, mechanistically, IAA-B in these mutants had distinct cellular etiology: ablation of Fn1 in using the Isl1^Cre^ knockin mice led to defective formation of the 4^th^ PAAs ^29^, while the ablation of Fn1 in the NC resulted in the regression of originally well-formed 4^th^ PAAs ^31^.

Integrins are a major class of transmembrane receptors that engage in signal transduction upon binding ECM proteins. Integrins are heterodimers of α and β chains. There are 18 α and 8 β subunits encoded by mammalian genomes, giving rise to 24 different αβ combinations ^32^. Integrin α5 complexes with integrin β1, forming the integrin α5β1 heterodimer ^33^. Integrin α5β1 binds the ECM glycoprotein fibronectin (Fn1), and regulates Fn1 assembly *in vivo* ^34^. Phenotypes resulting from either global or cell-type-specific ablations of integrin α5 (MGI gene symbol: Itga5) or Fn1 in mice are similar ^26, 28, 29, 31, 34-41^, indicating that integrin α5β1 is a major Fn1 signal transducer *in vivo*. Previously, we demonstrated that the expression of integrin α5β1 and Fn1 in the Isl1 lineages was required for the formation of the 4^th^ PAA and that the deletion of either integrin α5 or Fn1 in using the Isl1^Cre^ knockin strain resulted in IAA-B ^29^. To understand the mechanisms by which integrin α5β1 and Fn1 regulate AAA development, we analyzed SHF and endothelial cell dynamics in integrin α5^f/-^; Isl1^Cre^ and Fn1^f/-^; Isl1^Cre^ mutants during PAA formation and remodeling, spanning embryonic days (E) E9.5 - E11.5 of development. Our studies point to the essential roles of cell-ECM interactions mediated by integrin α5β1 and Fn1 at multiple stages of PAA formation and remodeling.

## Methods

### Animals

All experimental procedures were approved by the Institutional Animal Care and Use Committee of Rutgers University and conducted in accordance with the Federal guidelines for the humane care of animals.

### Tamoxifen injections

*Isl1*^*MerCreMer*^ knockin mice ^42^ and Mef2C-AHF-DreERT2 transgenic mice ^43^ were used for temporal labeling of vascular progenitors in the SHF. Tamoxifen was dissolved either in corn oil or in sesame oil at the concentration of 10 mg/ml. Labeling was done by i.p. injection of 300 μl of stock solution into pregnant females at multiple time points specified in the legend to Fig. 1. E0.5 was designated to be as noon on the day when the vaginal plug was found.

**Figure 1.**
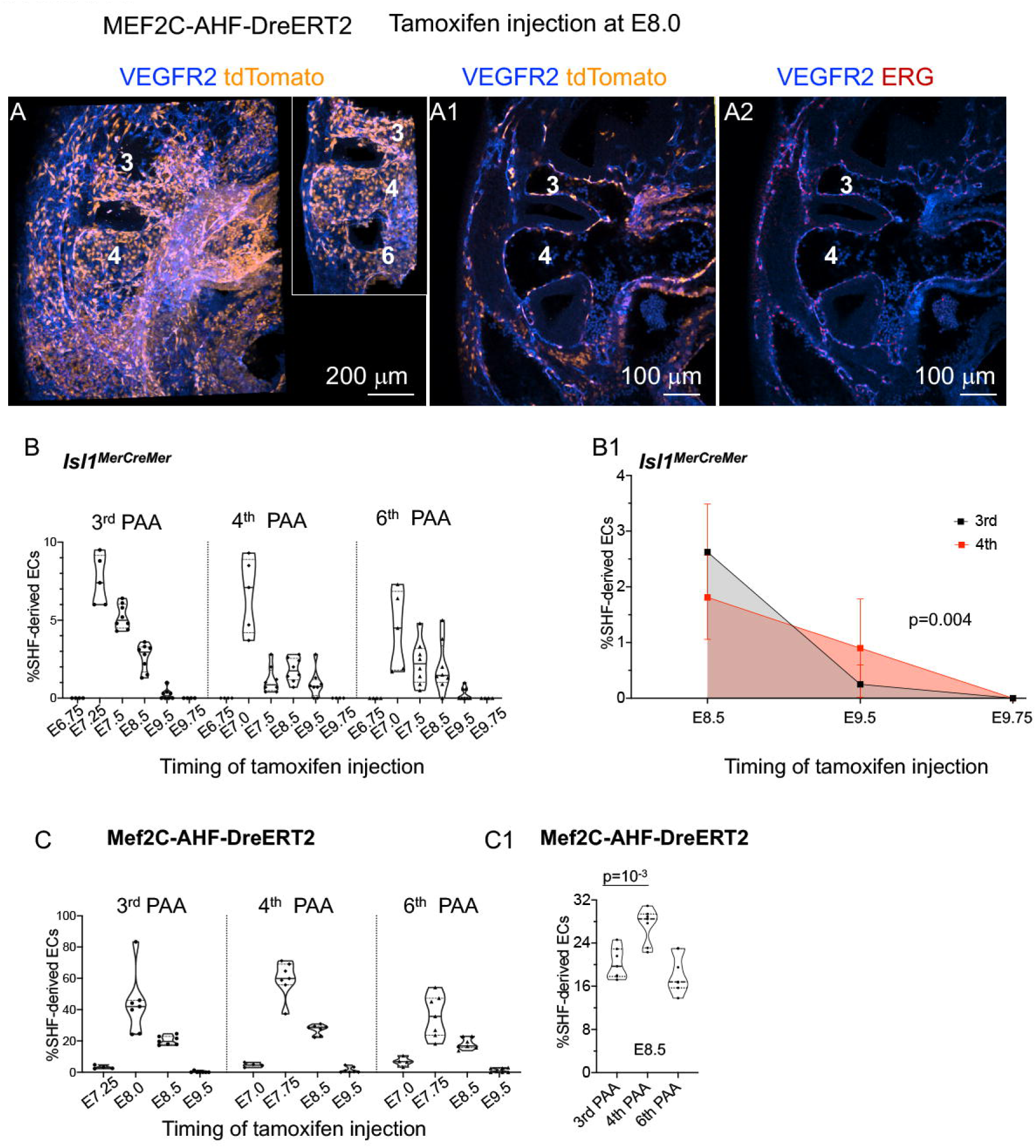
Endothelial PAA progenitors are present arise in the second heart field as early as E7.25. Isl1^MerCreMer^ and Mef2C-AHF-DreERT2 males were mated with the appropriate reporter females (see Methods). E0.5 was considered to be noon on the day of the vaginal plug. Tamoxifen was injected at specified times, embryos were dissected at E10.5, and stained to detect VEGFR2 (blue), ERG (red), or tdTomato (orange). **A.** Sagittal view and 3D reconstruction through the left pharyngeal region. Inset-3D reconstruction of PAAs. **A1 – A2.** Sagittal optical sections through the embryo shown in **A.** Labeling efficiency was quantified by calculating the ratio of the number of ERG^+^ tdTomato^+^ ECs to the total number of ERG^+^ ECs in PAAs using IMARIS spot function. **B.** Highest labeling of PAA endothelium occurred when tamoxifen was injected at E7.25 in Isl1^MerCreMer^ knockin mice. **B1.** SHF-derived cells continute to be added the 4^th^ PAA after E9.5. C. Peak labeling of PAA ECs occurred when tamoxifen was injected at E8.0 in Mef2C-AHF-DreERT2 strain. **C1.** Injection of tamoxifen at E8.5 led to a more efficient labeling of the 4^th^ PAAs than the 3rd and 6^th^ PAAs.

### Whole Mount Immunofluorescence staining

Labeling with BrdU, TUNEL, and staining with antibodies were performed as described ^29^, and analyzed using IMARIS (Bitplane, USA) ^9, 44^. Detailed procedures for staining, analysis and cell quantification is described in ^44^.

### Statistics

Statistical analyses were performed using Prism 8 software version 8.4.3. Specific statistical tests are indicated in figure legends.

## Results

### SHF contributes harbors PAA progenitors between E7.5 and E9.5

Previous work from our lab demonstrated that in the mouse, the majority of PAA endothelium is derived from the SHF, derived from either Mef2C-AHF-Cre- or Isl1^Cre^-expressing mesodermal lineages ^9^. To define the temporal window during which the SHF mesoderm harbors endothelial progenitors of the PAAs, we used *Isl1*^*MerCreMer*^ knockin mice ^42^ and Mef2C-AHF-DreERT2 transgenic mice ^43^ combined with pulses of tamoxifen to lineage-label the SHF mesoderm at different developmental times (Fig. 1). Tamoxifen was injected at discrete time points between E6.75 – E9.75, and embryos were dissected at E10.5 and stained to detect lineage labeling in the pharyngeal arches. Entire pharyngeal regions were imaged using confocal microscopy to quantify the contribution of lineage-labeled cells to the PAA endothelium (Fig. 1, panels A-A2). The expression of VEGFR2 and ERG were used to mark EC cell membrane and nuclei ^45^. The labeling of the cardiac outflow tract and the right ventricle using *Isl1*^*MerCreMer* /+^ mice was evident at all stages tested, indicating that our labeling technique was consistent with previous studies (data not shown) ^42^. Myocardial cells derived from the SHF are labeled when tamoxifen is injected as early as E6.5 in *Isl1*^*MerCreMer*^ knockin mice ^42^, however no PAA ECs were labeled when tamoxifen was injected at E6.75 in this strain (Fig. 1B) or at E7.25 in Mef2C-AHF-DreERT2 transgenic strain (Fig. 1C), suggesting that PAA EC progenitors arise later in the SHF relative to cardiomyocyte progenitors. The peak labeling of the PAA endothelium occurred when tamoxifen was injected at E7.25 in *Isl1*^*MerCreMer*^ strain (Fig. 1B) and at E8.0 in Mef2C-AHF-DreERT2 strain (Fig. 1C). While tamoxifen injection into *Isl1*^*MerCreMer*^ resulted in sparse labeling of PAA ECs, the injection of tamoxifen into Mef2C-AHF-DreERT2 transgenic mice led to the labeling of a much larger proportion of ECs in the PAAs (compare Fig. 1B with Fig. 1C). These differences likely reflect that the MerCreMer transgene is present as a single copy as it is knocked into the *Isl1* locus ^42^, while Mef2C-AHF-DreERT2 is a transgenic strain containing multiple copies of the Mef2C-AHF-DreERT2 transgene ^43^. In addition, the expression of *Is1* is downregulated commensurate with endothelial differentiation ^46^. Thus, potentially low levels of *MerCreMer* expression in EC precursors could have resulted in low labeling of endothelial progenitors in *Isl1*^*MerCreMer*^ mice relative to Mef2C-AHF-DreERT2 strain. The difference in the timing of peak EC labeling in the PAAs between *Isl1*^*MerCreMer*^ and Mef2C-AHF-DreERT2 strains is likely due to the earlier onset of *Isl1* expression compared with the expression of the Mef2C-AHF-DreERT2 transgene; in fact, Isl1 regulates the expression of Mef2C and the activation of the Mef2C-AHF enhancer ^47, 48^. Correspondingly, our experiments demonstrate that the peak endothelial labeling of PAAs in *Isl1*^*MerCreMer*^ strain precedes that of Mef2C-Dre-ERT2 strain by 18 hours (compare Fig. 1B with Fig. 1C). Interestingly, the accrual of SHF-derived ECs into the 4^th^ arch continues past E8.5 as more SHF-derived cells are labeled in the 4^th^ PAAs than in the 3^rd^ and 6^th^ when tamoxifen is injected at E8.5 and E9.5 (Fig. 1B1, 1C1). Thus, our labeling experiments show that the SHF mesoderm harbors PAA endothelial progenitors between approximately E7.5 and E9.5 of embryonic development.

To analyze the contribution of the SHF to the PAA endothelium quantitatively and to compare two mouse strains commonly used to label the SHF, we imaged the entire pharyngeal arch region corresponding to arches 3, 4, and 6, and quantified the proportion of SHF-lineage labeled ECs in the PAAs from E10.5 embryos derived from Isl1^Cre^ knockin and Mef2C-AHF-Cre transgenic lines (Fig. 2). The majority of SHF-derived cells in the pharyngeal arches 3 – 6 are found in the endothelium at 37somites, as seen in sections through the pharyngeal arch region (Fig. 2A – B). Each PAA is comprised of a similar number of ECs (Fig. 2C). However, there were differences in PAA labeling among embryos isolated from Isl1^Cre^ and Mef2C-AHF-Cre mice (Fig. 2D, E). In the constitutive *Isl1*^*Cre*^ knockin strain, the SHF contribution to the 3^rd^ and 4^th^ PAA endothelium was 79±6% and 77±10%, respectively, and 57±12% to the 6^th^ PAA (Fig. 2D). While the SHF contribution to the 3^rd^ PAA was 45±8% in Mef2C-AHF-Cre transgenic line, which is significantly different from *Isl1*^*Cre*^ knockin strain (p<10^−5^, one-way ANOVA with Tukey’s correction for multiple testing). The SHF contribution to the PAA endothelium of the 4^th^ and 6^th^ PAAs were similar between the two strains (p>0.2, one-way ANOVA with Tukey’s correction for multiple testing). The difference in the contribution of the SHF to the 3^rd^ PAA between the two strains likely reflects the earlier onset of Cre expression in the *Isl1*^*Cre*^ knockin strain relative to Mef2C-AHF-Cre transgenic line ^47^. These data suggest that about half of the 3^rd^ PAA progenitors arise and leave the SHF prior to the activation of Mef2C-AHF-Cre. These data also indicate that the deletion of one Isl1 allele, as in the *Isl1*^*Cre*^ knockin strain, does not impair the contribution of the SHF to the PAA endothelium. In summary, our data show that the 4^th^ PAAs differ from the 3^rd^ and the 6^th^ PAAs in the timing during which SHF cells are added, and differ from the 6^th^ PAA in the proportion of SHF-derived cells.

**Figure 2.**
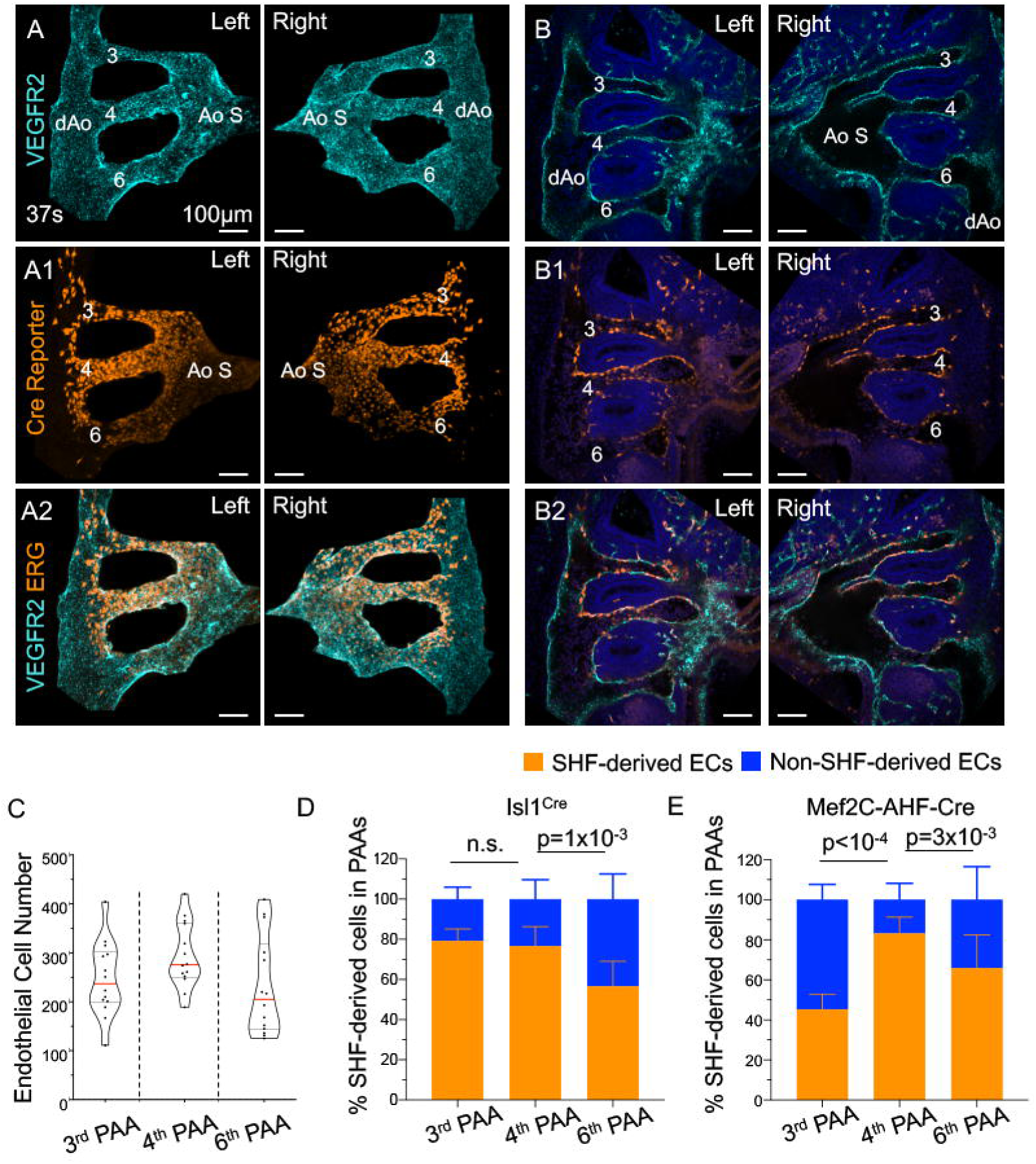
Majority of PAA ECs are SHF-derived; there are differences in the contribution of the SHF to the PAA endothelium depending on the strain used. Mef2C-AHF-Cre; ROSA^tdTomato^ embryos (35 – 37 somites) were stained with antibodies to VEGFR2 (turquoise) to detect endothelial cells, tdTomato (orange) to detect SHF-derived cells, and DAPI (blue) to stain cell nuclei. **A – A2.** 3D reconstructions of PAAs and their connections with the dorsal aorta (dAo) and the aortic sac (Ao S). **B – B2.** Sagittal optical sections to show the distribution of all SHF-derived cells in the pharyngeal arches. PAAs 3 – 6 are labeled. Magnifications are the same in all panels. All scale bars are 100 μm. **C.** The number of VEGFR2^+^EGR^+^ cells in the pharyngeal arches was quantified in 5 E10.5 embryos at 35 – 37 somites using IMARIS. Each dot is one arch. Red line marks the median. Black lines mark quartiles. Differences among the three PAA pairs are not significant, p>0.1 by one-way ANOVA with Tukey’s correction for multiple testing. **D – E.** The percentage of VEGFR2^+^EGR^+^ cells expressing the Cre reporter was determined in each PAA (orange bars). Blue bars are the percent of VEGFR2^+^EGR^+^ cells that were not labeled with the Cre reporter. **D.** The use of constitutive Isl1^Cre^ strain resulted in labeling of more than 80% of ECs in the 3^rd^ and 4^th^ PAAs. **E.** The use of Mef2C-AHF-Cre strain resulted in a significantly higher labeling of the 4^th^ PAAs than the 3^rd^ and the 6^th^. The difference in the labeling efficiency of 6^th^ PAAs between the two strains was not significant, p> 0.2. All statistical analyses were performed using one-way ANOVA with Tukey’s correction for multiple testing.

### Cell-ECM interactions mediated by integrin α5β1 specifically regulate the accrual of SHF-derived cells into pharyngeal region

Studies described above together with our previous work ^9^ have established a framework for the analyses of EC dynamics and their genetic regulation during the morphogenesis of AAA and its major branches. Our previous studies demonstrated that the deletion of either integrin α5 or Fn1 in the Isl1 lineages resulted in the defective formation of the 4^th^ PAAs at E10.5, and consequently, IAA-B and retro-esophageal right subclavian artery (RERSA), in these mutants ^29^. IAA-B and RERSA are anomalies resulting from defective morphogenesis of the left and right 4^th^ PAAs, respectively ^1, 3^. To determine the mechanisms by which integrin α5β1 and Fn1 regulate the formation of the 4^th^ PAAs, we analyzed PAA development at distinct stages of embryonic development using whole-mount immunofluorescence followed by quantitative analyses of SHF-derived populations and their dynamics.

PAAs form through the coalescence of pharyngeal arch EC plexus, a network of small blood vessels ^9, 10^. All pharyngeal arch ECs are located within the plexus at E9.5. At E10.5 (33 – 34 somite stage), 50% of the 4^th^ arch endothelium is found within the PAA (the vessel surfaced in green in (Fig. 3B, C) and 50% is in the plexus (pink in Fig. 3B, C)^9^. About 50% of integrin α5^f/-^; Isl1^Cre^ mutants have defective 4^th^ PAAs, and consequently, 50% of these mutants develop IAA-B and RERSA ^29^. We found that 4^th^ PAA is absent in 50% of mutants at 32 – 34 somites (Fig. 3D – F). Instead, the endothelium in the 4^th^ arches is in the form of a plexus of small blood vessels (marked in pink in Fig. 3E, F). A small 4^th^ PAA eventually formed in these mutants by 36 – 39 somites (Fig. 3J, marked in green in Fig. 3K, L; compare with the 4^th^ PAA surfaced in green in control Fig. 3G-I). Similarly, the formation of the 4^th^ PAA was delayed in Fn1^f/-^; Isl1^Cre/+^ mutants (Sup. Fig. 1). This defect was specific to the 4^th^ PAA, as the 3^rd^ and 6^th^ PAAs formed normally in the mutants (vessels surfaced in white and red in Fig. 3). The incidence of IAA-B and RERSA is 50% in integrin α5^f/-^; Isl1^Cre^ and Fn1^f/-^; Isl1^Cre/+^ mutants, which is the same as the incidence of defective 4^th^ PAAs at E10.5 ^29^.

**Figure 3.**
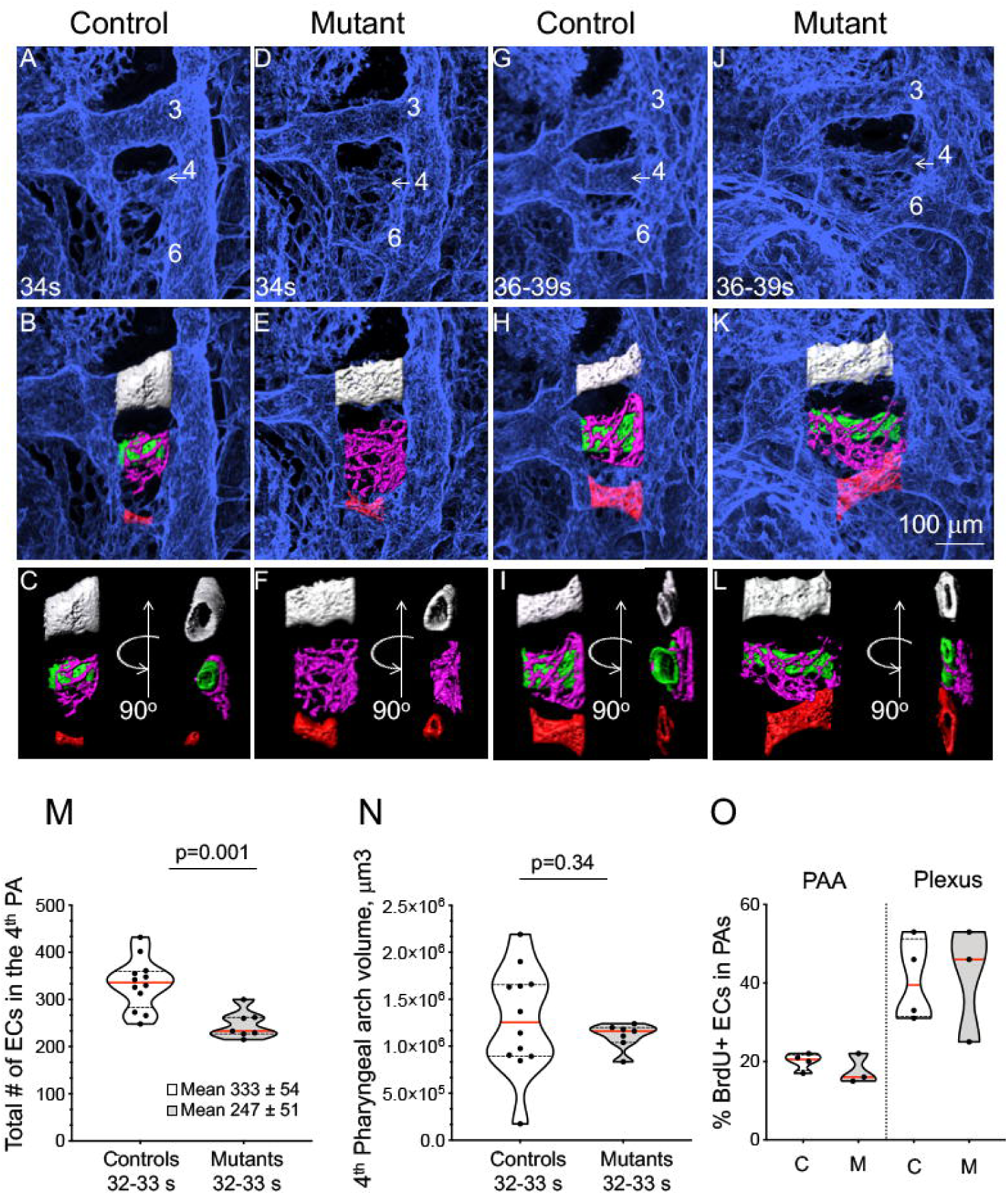
Formation of the 4^th^ PAA is delayed in integrin α5^flox/-^; Isl1^*Cre*^ mutants. α5^flox/+^; Isl1^*Cre*^ control and α5^flox/-^; Isl1^*Cre*^ mutant embryos were dissected at different somite stages at E10.5 and stained to detect Pecam1. PAAs are numbered and somite stages are indicated in the first row. **A, D, G, J.** 3D reconstructions of whole-mount Pecam 1 staining (light blue). **B, E, H, K.** PAA endothelium in the 3^rd^, 4^th^ and 6^th^ arches shown in the row above was surface-rendered white, green and red, respectively. In addition, the plexus endothelium in the 4^th^ arch was surface-rendered in pink. **C, F, I, L.** Left side and ventral views of surface-rendered PAAs and the plexus. Development of the 4^th^ PAAs was specifically affected in the mutants (**E, F**). Magnification is the same in all panels. Scale bar is 100 μm. **M.** Total number of endothelial cells was quantified as described in Methods. Mutants have EC deficiency in the 4^th^ arch as 32 – 33 somites. **N.** The sizes of the 4^th^ arches are comparable between controls and mutants. **O.** EC proliferation in the PAA and the plexus in the 4^th^ arches was similar in controls (C) and mutants (M). In all plots, solid lines mark the median, dashed lines mark the quartiles. Each dot marks one arch. At least 3 mutants and 3 controls were assayed. Statistics were evaluated using 2-tailed, unpaired Student’s t test with Welch’s correction for unequal standard deviation between samples.

Therefore, we further investigated the mechanisms by which integrin α5β1 and Fn1 regulate the formation of the 4^th^ PAAs.

We hypothesized that the defective formation of the 4^th^ PAAs in our mutants could be due to insufficient EC numbers, defective EC proliferation, or survival. To test these hypotheses, we evaluated total EC numbers in the 4^th^ pharyngeal arches of controls and mutants. To quantify EC number, we stained E10.5 embryos with the antibodies to ERG, a transcription factor enriched in the endothelia and either VEGFR2 or Pecam 1, expressed on EC surface, as described ^44^. These experiments showed that integrin α5^f/-^; Isl1^Cre^ and Fn1^f/-^; Isl1^Cre/+^ mutants had decreased total number of ECs in the 4^th^ arches at 32 – 33 somites relative to controls (Fig. 3M and Sup. Fig. 1A-C). Despite this decrease in EC numbers, the size of the 4^th^ arches, the tissues within which PAAs form, was not affected (Fig. 3N). EC proliferation in the 4^th^ arch was also not affected in the mutants (Fig. 3O), and neither was cell survival (Sup. Fig. 2). Thus, EC deficiency in the 4^th^ pharyngeal arches of the mutants was not due to defective cell proliferation or survival. The majority of VEGFR2^+^ cells in the pharyngeal region of E9.5 embryos were labeled with GFP (Sup. Fig. 3A – A2, B – B2), indicating that SHF cells in the pharyngeal region of the mutants were not impaired in the acquisition of EC fate. VEGFR2 is expressed the 4^th^ pharyngeal arch endothelium at E9.5, which is a day earlier than Pecam 1 ^10^. To determine whether the maturation of pharyngeal arch ECs was affected in the mutants at E10.5, we co-stained embryos with antibodies to VEGFR2 and Pecam 1. Despite defective 4^th^ PAA formation, all VEGFR2^+^ cells in pharyngeal arches of the mutants also expressed Pecam1 at E10.5 (Sup. Fig. 3C, D, C1, D1), ruling out maturation as a cause for decreased EC numbers in the 4^th^ pharyngeal arch.

Since the maturation, proliferation, and survival of ECs were not affected in our mutants, we tested the hypothesis that defective recruitment of progenitor cells into the pharyngeal arches was the cause for decreased EC numbers in the 4^th^ arch. As we established before, the majority of PAA endothelium arises from the SHF (Fig. 2D, E and ^9^, and the accrual of SHF-derived cells into the pharyngeal arches is mostly complete by E9.5 (Fig. 1B-C). To quantify the number of SHF-derived cells in the pharyngeal mesenchyme, we used ROSA^nT-nG^ reporter mice, in which nuclear localization sequences were fused with tdTomato and EGFP, leading to the expression nuclear-localized EGFP upon Cre-induced recombination. We found that the deletion of integrin α5 in the Isl1 lineages impaired the accrual of SHF cells into the pharyngeal region (Fig. 4A-B), while the accrual of SHF-derived cells into the heart was not affected (Fig. 4C).

**Figure 4.**
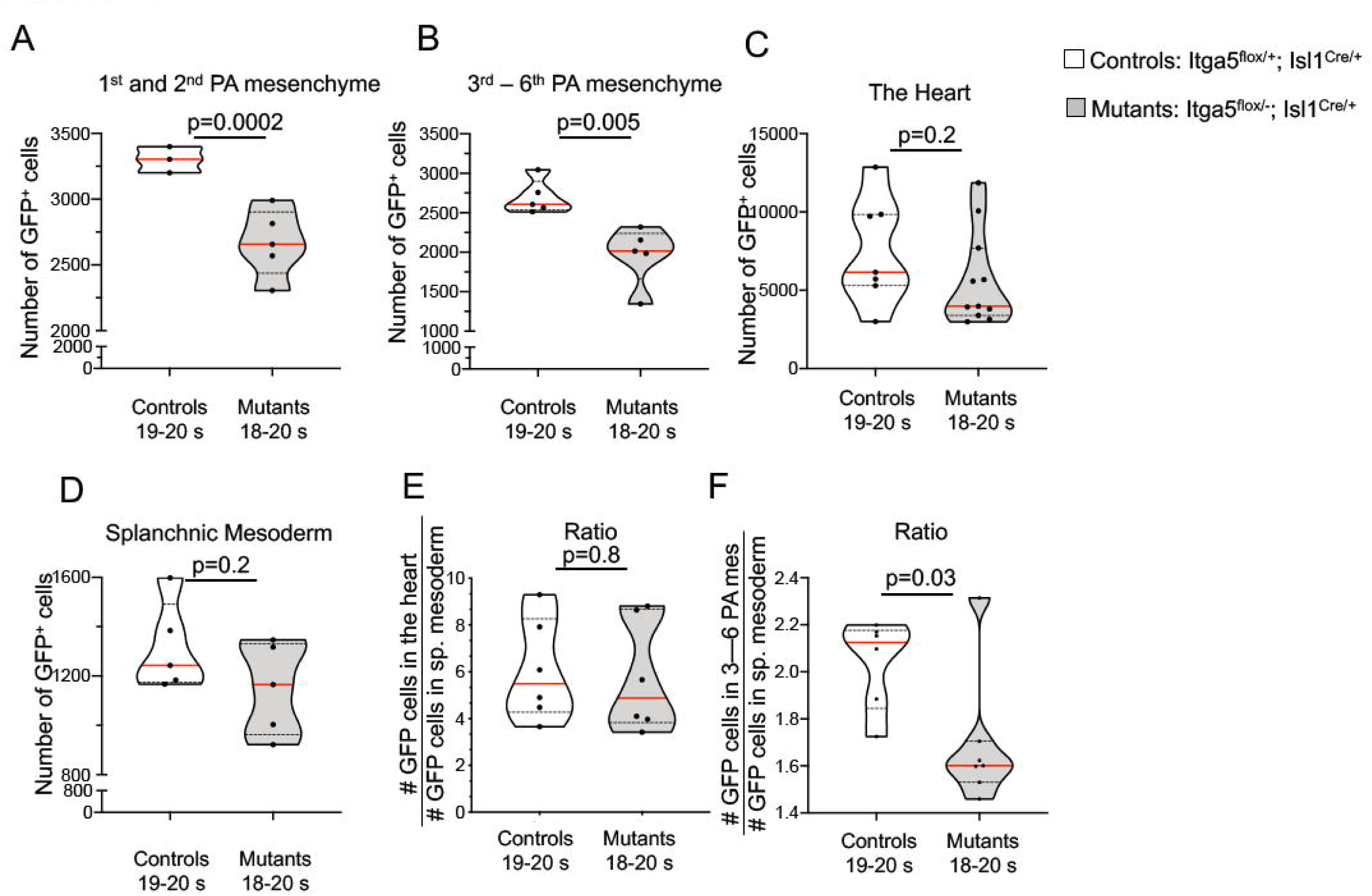
The expression of integrin α5 in the Isl1 lineages is required for the accrual of SHF-derived cells to the pharyngeal mesenchyme. Control and mutant embryos carrying one ROSA^nTnG^ reporter allele were dissected at E9.5 (18 – 20 somite stage) and stained with DAPI and anti-GFP antibodies. Whole embryos were imaged and the number of GFP^+^ cells (SHF-derivatives) in the arch mesenchyme, splanchnic mesoderm, and in the heart was quantified as described in Sup. Fig. 4. **A.** The total number of SHF cells in the mesenchyme of the 1^st^ and 2^nd^ arches was decreased in the mutants. **B.** The total number of of GFP^+^ cells in the pharyngeal mesenchyme corresponding with the future arches 3 – 6 was decreased in the mutants. The number of SHF cells in heart (**C**) and the splanchnic mesoderm (**D**) was not affected. **E.** The proportion of GFP^+^ cells in the heart relative to GFP^+^ cells in splanchnic mesoderm was not affected in the mutants. **F.** The proportion of GFP^+^ cells in the posterior pharyngeal mesenchyme relative to the number of GFP^+^ cells in splanchnic mesoderm was significantly decreased in the mutants. Each dot marks one embryo, red lines mark medians, dotted lines mark quartiles; p values were determined using unpaired, 2-tailed Student’s t tests.

Splanchnic mesoderm within the dorsal pericardial wall harbors both cardiac and vascular progenitors. To test whether the deficiency in the pharyngeal SHF-derived mesoderm was due to the decrease in SHF cells numbers in the splanchnic mesoderm, we used IMARIS to surface cells within this region and quantified the number of GFP^+^ cells (see Sup. Fig. 4 for details on surfacing). These experiments showed that the number of GFP^+^ cells in the splanchnic mesoderm within the dorsal pericardial wall was similar between controls and mutants (Fig. 4D). Next, we computed the proportion of GFP^+^ cells in the pharyngeal mesenchyme or in the heart relative to the number of GFP^+^ cells in the splanchnic mesoderm. While the latter ratio was not affected in the mutants (Fig. 4E), the former was significantly decreased in the mutants (Fig. 4F), suggesting that there is a defect either in the specification of pharyngeal progenitors in the SHF or in their exit from the SHF. Taken together, these experiments indicate that ECM microenvironment sensed through the signaling by integrin α5β1 is important for the accrual of the SHF-derived mesoderm to the pharyngeal arches (Model in Fig. 8, PAA formation panel A1).

### Integrin α5β1 and fibronectin regulate the remodeling of pharyngeal plexus into the 4^th^ PAAs independently of EC numbers

Interestingly, the number of SHF-derived cells and ECs in the mutants recovered by the 34-35 somite stage, and was similar to that of controls (Fig. 5A). Total number of GFP^+^ cells also recovered (Fig. 5B). The percentage of GFP^+^ ECs in the pharyngeal arches of controls and mutants were comparable (Fig. 5C), indicating that the recovery was not due to the recruitment of ECs from an alternative mesodermal source. The recovery of pharyngeal EC numbers was likely mediated through the proliferation of SHF-derived ECs. The basis for this conclusion is the following. The proliferation index of pharyngeal arch ECs was unaltered in the mutants (Fig. 3O), and the proliferation index of ECs in the pharyngeal plexus is 2-fold higher than that of PAA ECs, both in controls and in mutants (Fig. 3O, plexus). Since the proportion of ECs in the pharyngeal plexus is higher in the mutants than in controls (Sup. Fig 1D), the higher proliferation index of plexus ECsin the mutants is likely responsible for EC recovery. Our quantitative analyses indicate that PAA formation phenotypes in integrin α5^f/-^; Isl1^Cre/+^ and Fn1^f/-^; Isl1^Cre/+^ mutants are indistinguishable from one another (Sup. Fig. 1), suggesting that integrin α5β1 is a major receptor transducing Fn1 signals within the pharyngeal microenvironment.

**Figure 5.**
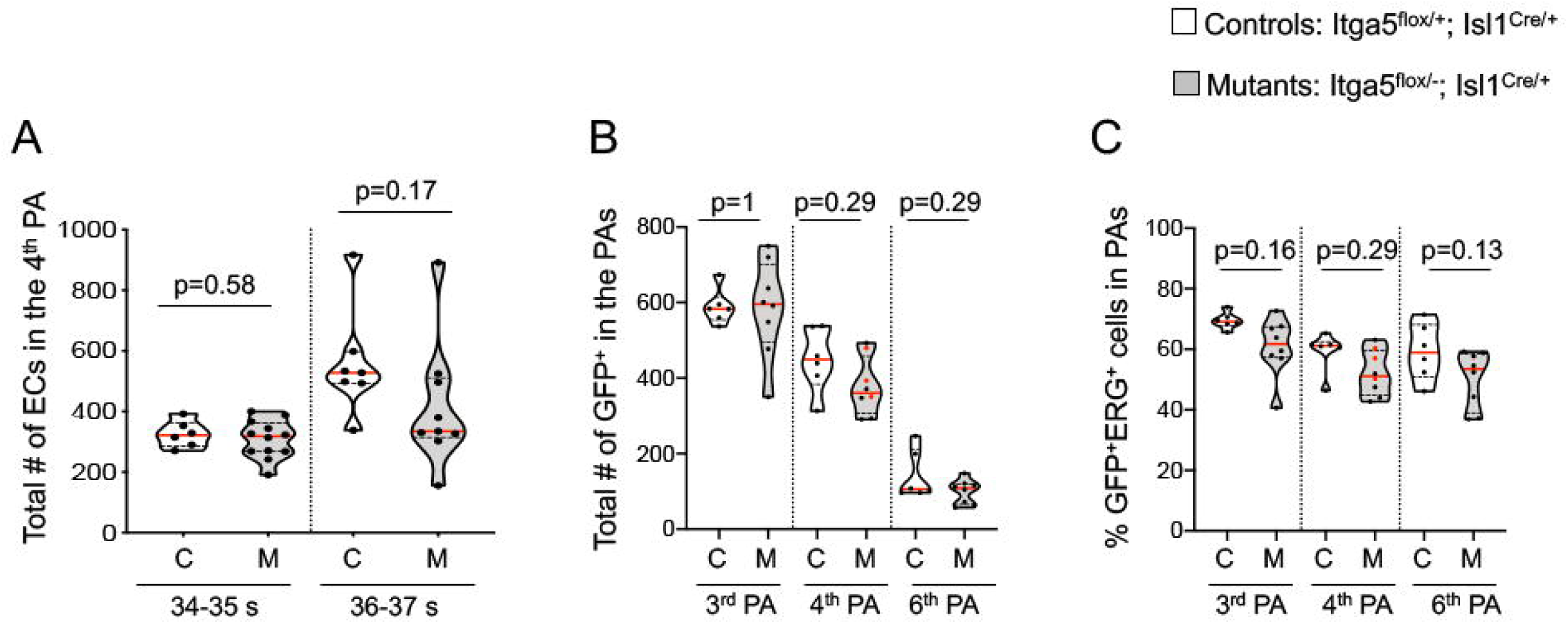
Recovery of EC numbers in integrin α5^flox/-^; Isl1^Cre/+^ mutants. **A.** Total EC number has recovered in integrin a5^flox/-^; Isl1^Cre/+^ mutants by the 34^th^ somite stage. **B.** Total number of SHF-derived mesodermal cells has recovered in the pharyngeal arches in integrin α5^flox/-^; Isl1^Cre/+^ mutants by the 34^th^ somite stage. **C.** The fraction of SHF-derived ECs in pharyngeal arches is comparable among control and mutant embryos. This fraction was calculated by quantifying the number of GFP^+^ERG^+^ cells and dividing by the total number of ERG^+^ cells in the entire pharyngeal arches (e.g. ECs in PAA and plexus were quantified). Statistical significance was evaluated using one-way ANOVA with Tukey’s correction for multiple testing.

Despite the recovery of EC numbers (Fig. 5A), PAAs remained thin in integrin α5^f/-^; Isl1^Cre^ and Fn1^f/-^; Isl1^Cre/+^ mutants (Fig. 3G – L), and there was 2 – 3-fold decrease in the proportion of pharyngeal arch ECs in the 4^th^ PAAs at all stages analyzed at E10.5 (Fig. 6A). The size of the 4^th^ PAA increases between 32 – 39 somites, as more ECs are added to the PAA from the plexus (Sup. Fig. 5), and is reflected in the percent of pharyngeal arch ECs in the PAA ^9^. In controls, plexus ECs in the 4^th^ arch begin coalescing into the PAA when embryos reach between 31 and 32 somites ^9^ (Sup. Fig. 5). These rearrangements result in an initially thin 4^th^ PAAs, in which approximately 50% of the pharyngeal arch ECs are in the plexus and 50% in the PAA at 32 – 34 somite stage ^9^. As the development proceeds, by 36-39 somite stage, > 60% of the 4^th^ pharyngeal arch endothelium becomes incorporated into the 4^th^ PAA ^9^. Thus, the percentage of the pharyngeal arch endothelium in the 4^th^ PAA can be taken as a measure of PAA formation. The higher the proportion, the larger the PAA ^9^.

**Figure 6.**
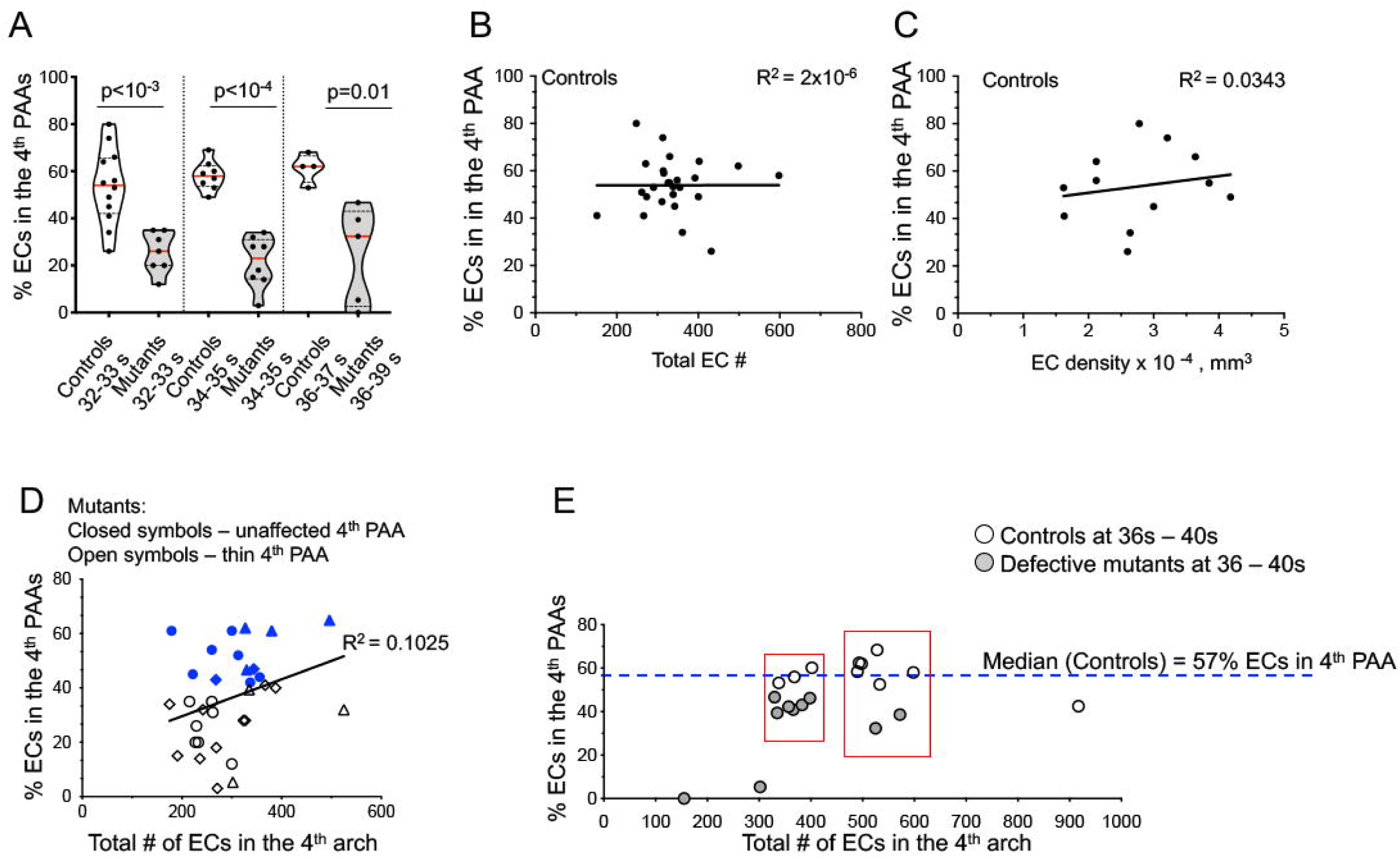
Integrin α5β1 and Fn1 regulate the remodeling of EC plexus during the formation of the 4^th^ pharyngeal arch arteries. **A.** The proportion of ECs in the 4^th^ PAAs in the mutant is significantly lower than in controls at all stages analyzed at E10.5, including the stages when the EC population in the 4^th^ pharyngeal arch has recovered in the mutants; 2-tailed, unpaired Student’s t test. **B – C.** Linear regression analyses indicate the absence of linear correlation between the size of the 4^th^ PAA and EC number (**B**) or density (**C**). PAA size is expressed as the percentage of pharyngeal arch endothelial cells in the 4^th^ PAA in control embryos on the y-axis. **D.** Total EC number (x-axis) from mutants with defective (open symbols) or unaffected 4^th^ PAA (closed symbols) were plotted against the size of the 4^th^ PAAs, y-axis. Regression analysis indicated low correlation between these properties. Circles: 32 – 33 somite embryos, rhombi: 34 – 35 somite embryos, triangles: 36 – 39 somite embryos. **E.** The rearrangement of the endothelial plexus into the 4^th^ PAAs is defective in mutants relative to controls with the same number of endothelial cells in the 4^th^ arch (red boxes). EC – endothelial cell(s). Controls: α5^f/+^; Isl1^Cre/+^ and Fn1^f/+^; Isl1^Cre/+^ embryos; Mutants: α5^f/-^; Isl1^Cre/+^ and Fn1^f/-^; Isl1^Cre/+^ embryos.

To understand the mechanisms by which integrin α5β1 and Fn1 regulate the remodeling of the uniform endothelial plexus into the PAA in the 4^th^ arch, we examined EC dynamics in control and mutant embryos at three time points, corresponding with 32 – 33 somites, 34 – 35 somites, and 36 – 39 somites. These stages span about 6 hours on the 10^th^ day of mouse embryonic development. The formation of the 4^th^ PAAs lagged in mutants relative to controls at all time points tested during E10.5 (Fig. 6A and Sup Fig. 1D, E), and 7 of the 16 embryos analyzed contained only a plexus of ECs and lacked the 4^th^ PAAs at the 32 – 34 somite stage (Sup. Fig. 1E), a stage at which over 50% of the 4^th^ arch endothelium in controls is located within the 4^th^ PAAs ^9^ (Fig. 6A and Sup Fig. 1E).

Since mutant embryos had fewer ECs in the 4^th^ pharyngeal arches compared with controls prior to the 36^th^ somite stage, we performed correlation analyses to test the hypothesis that the formation of the 4^th^ PAAs depended on the total EC number or EC density in the 4^th^ pharyngeal arches. As described above, the percentage of pharyngeal arch ECs in the PAA relative to the plexus can be taken as a measure of PAA formation (Sup. Fig. 5) ^9^. Thus, for these analyses, we quantified EC numbers in control embryos isolated between 32 to 39 somite stages and plotted them against the percent of ECs in the 4^th^ PAAs (Fig. 6B). Despite the sharp, over a 3-fold increase in the number of ECs in the 4^th^ arches between these stages ^9^, the formation of the 4^th^ PAAs was independent of the total EC number in the 4^th^ pharyngeal arch tissue (Fig. 6B) or EC density (Fig. 6C) in controls throughout the 10^th^ day of embryonic development. Similarly, correlation analysis of PAA formation in the mutants with defective (thin) and normal 4^th^ PAAs, showed that similar to controls, the rearrangement of plexus ECs into the PAA did not depend on the number of total number of ECs in the 4^th^ pharyngeal arches of mutants (Fig. 6D).

Next, we compared PAA formation in controls and mutants that had a similar number of ECs in the 4^th^ pharyngeal arches (Fig. 6E). These analyses showed that in groups of mutant and control embryos with a similar number of ECs, the percent of ECs in the PAAs was lower in the mutants (boxes in Fig. 6E). These data indicate that the reorganization of the plexus ECs into the PAA in the 4^th^ pharyngeal arch does not depend on the EC number at E10.5, and is regulated by factors extrinsic to the pharyngeal arch endothelium. In summary, our studies indicate that during the 10^th^ day of embryonic development, cell – ECM interactions mediated by integrin α5β1 and Fn1 are essential for the remodeling of the initially uniform vascular plexus into the PAA in the 4^th^ pharyngeal arches (Model in Fig. 8, PAA formation panel A2).

### The expression of integrin α5 in the Isl1 lineage is required for the differentiation of neural crest cells into vascular smooth muscle cells

In the Tbx1^+/-^ mouse model of 22q11 deletion syndrome, PAA formation recovers in 50 – 68% of the mutant mice ^49, 50^. To determine whether the rearrangement of the endothelial plexus in the 4^th^ arch was blocked or delayed, we examined E11.5 embryos. The incidence of IAA-B and RERSA in integrin α5^f/-^; Isl1^Cre^ mutants is 50%, which is the same as the incidence of defective 4^th^ PAA formation. Therefore, we expected to find absent or thin 4^th^ PAAs in the mutants at E11.5. Contrary to our expectations, the formation of the 4^th^ PAAs recovered in the mutants by E11.5, and PAA perimeters in the mutants were comparable with controls (Fig. 7A, n=8). Consistent with the recovery of SHF-derived ECs numbers by 33 – 35s, PAA ECs were GFP^+^ cells in the mutants as in controls (compare Fig. 7C1 with 7D1, arrowheads). Regression of left 4^th^ PAAs results in IAA-B and regression of the right 4^th^ results in RERSA ^51, 52^. Since 50% of *integrin* α*5*^*flox/-*^; *Isl1*^*Cre*^ mutants develop 4^th^ arch artery defects, such as IAA-B and RERSA ^29^, these data indicated that the 4^th^ PAAs eventually regress in the mutants. Arch artery regression is commonly caused by the defective differentiation of neural crest cells surrounding the PAA endothelium into vascular smooth muscle cells, VSMCs ^52-57^. In pharyngeal arches, VSMCs exclusively arise from NC-derived cells ^49, 58, 59^. To determine whether the differentiation of NC-derived cells into VSMCs was affected in our mutants, we analyzed VSMC differentiation in the pharyngeal arches. For these experiments, we calculated the fraction of vessel perimeter covered by alpha smooth muscle actin (αSMA)-expressing cells, using previously-developed methodology ^31^. We found that the differentiation of NC-derived cells into VSMCs was severely diminished around the PAAs in the mutants (quantified in Fig. 7B; compare sections in Fig. 7C, D, magnified in Fig. 7C2, D2; zoom-out panels are in Sup. Fig. 6). The decrease in αSMA expression was not due to NC cell death (Sup. Fig. 2).

**Figure 7.**
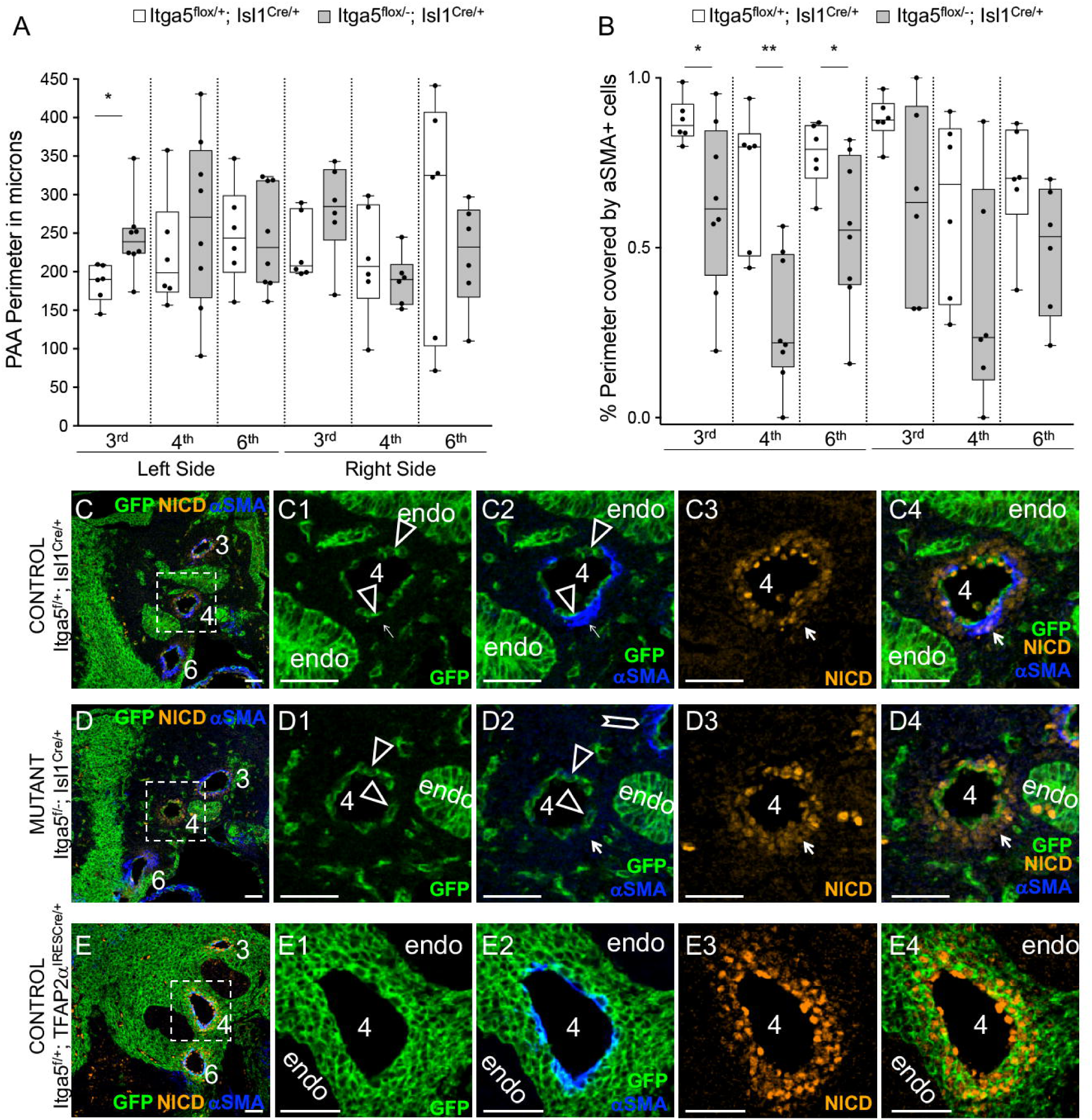
The expression of integrin α5β1 in the Isl1 lineages regulates the differentiation of neural crest-derived cells into VSCMs at E11.5. **A.** PAA perimeter has recovered in size in the mutants by E11.5. **B.** Smooth muscle coverage of the left 4^th^ and left 6^th^ PAA was deficient in the mutants. **C – D.** Despite defective differentiation of NC cells into VSMCs, the activation of Notch in the pharyngeal arch mesenchyme was not altered in the mutants. PAAs are numbered. PAA lumens at E11.5 are derived from the Isl1^Cre^ lineage (green, arrowheads in **C1** and **D1**). αSMA^+^ cells are GFP-negative in Isl1^Cre^ strain (arrows in **C1-C2**). **C2 – D2.** VSMC differentiation assayed by the expression of alpha smooth muscle actin (αSMA, blue) is specifically affected around the 4^th^ PAAs in the mutants (compare regions marked by the arrows in **C2** and **D2**). The activation of Notch assayed by the expression of NICD is not altered in the mutants with defective VSCM differentiation (arrows in **C3, C4** and **D3, D4**). **E.** Fate map using TFAP2α^IRESCre^ shows the location of NC-derived cells in the pharyngeal arches. Note extensive colonization of the mesenchyme between the endodermal pouches (endo) by the TFAP2α^IRESCre^ lineage. **E2.** αSMA^+^ cells are GFP^+^ in TFAP2α^IRESCre^ strain. All scale bars are 50 μm. Additional zoom-out views are in Sup. Fig. 6.

Even though, the *Isl1* lineage marks a subset NC-derived cells ^60^, Isl1 protein is not expressed in NC-lineage cells in the pharyngeal arches, and Isl1 lineage does not label cells adjacent to the PAA endothelium (Fig. 7C1, D1) ^29^. Moreover, comparison of NC lineage (Fig. 7E – E4) and Isl1 lineage maps at E11.5 demonstrates that αSMA expression coincides with the NC lineage (Fig. 7E2), but not with Isl1 lineage-labeled cells (arrows in Fig. 7C1 and C2 point to αSMA-expressing cells; arrowheads point to GFP^+^ PAA endothelium). These studies indicate that the expression of integrin α5 in the Isl1 lineage(s) regulates the differentiation of NC cells into VSMCs in a non-cell autonomous manner. These results are consistent with our previous experiments demonstrating that the expression of integrin α5 in the *Mesp1* lineage marking the anterior mesoderm regulates the differentiation of NC cells into VSMCs around the 4^th^ PAA ^59^. Since the deletion of integrin α5 in the *Mesp1* lineage does not result in defective or delayed PAA formation ^59^, these data taken together, indicate that the defect in VSMC differentiation in *integrin* α*5*^*flox/-*^; *Isl1*^*Cre*^ mutants is not merely due to the delayed accrual of arch artery endothelium, or delayed remodeling of the vascular plexus into the PAA. In summary, our studies also indicate that in addition to regulating 4^th^ PAA formation, integrin α5 expressed in the *Isl1* lineages plays an independent role in arch artery morphogenesis, namely in the differentiation of NC-derived cells into VSMCs (Fig. 8A).

**Figure 8.**
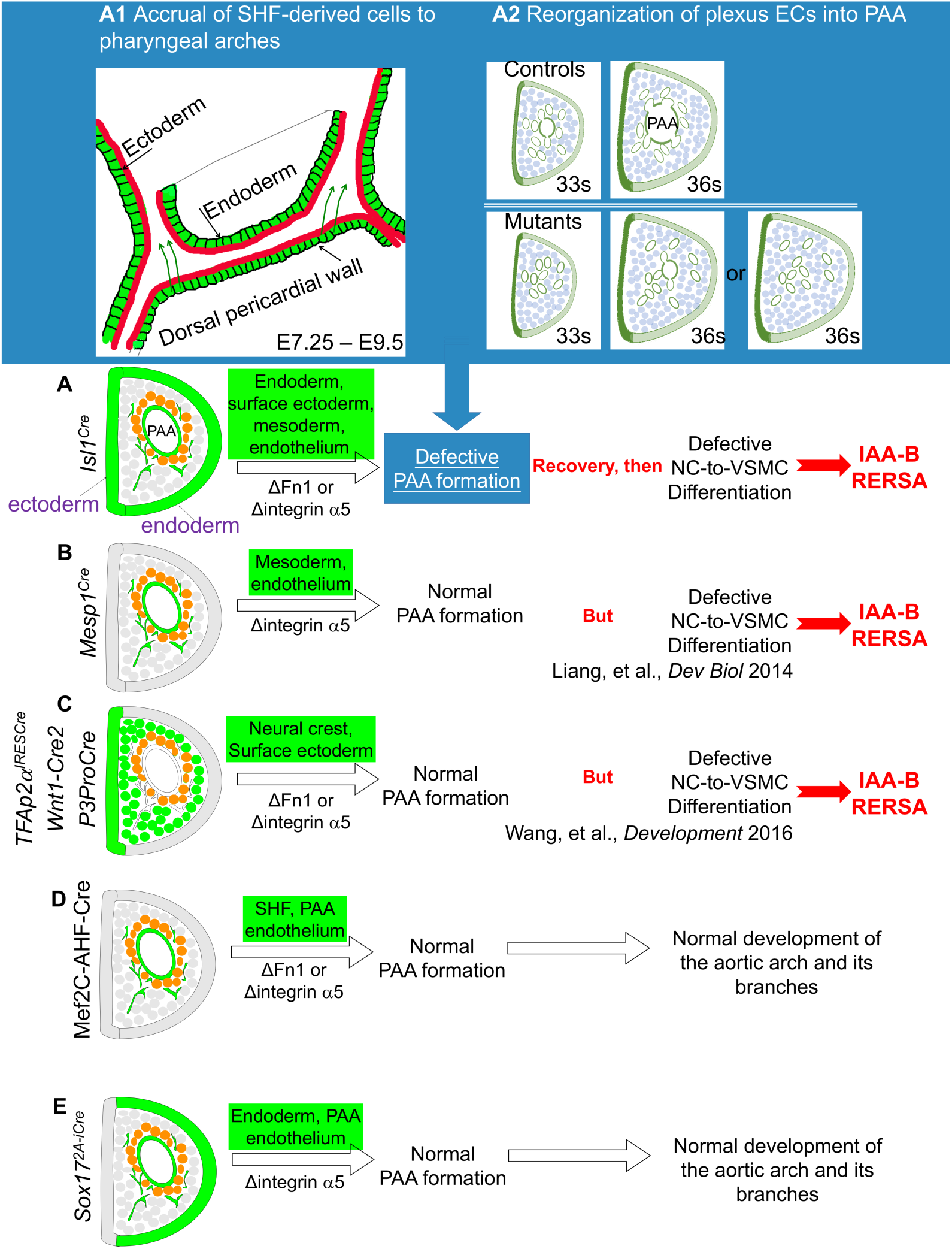
Cell – ECM interactions play essential roles at multiple stages during the development of the aortic arch and its branches. **A – E.** Depictions of coronal sections through the 4^th^ pharyngeal arch at E10.5. Lineages are marked in green, NC-derved cells are depicted as circles, NC-derived cells next to the PAA endothelium are marked in orange. **A.** The expression of integrin α5β1 and Fn1 in the Isl1 lineages is required for the formation of the 4^th^ PAAs (see **A1** and **A2** in the blue inset) as well as for NC-to-VSMC differentiation. The deletion of integrin α5 or Fn1 in the Isl1 lineages leads to IAA-B and RERSA. **A1 – A2.** Stages during which integrin a5 and Fn1 regulate the 4^th^ PAA formation. **A1.** A schematic of Isl1 lineages (green) in the pharynx at E8.5 – E9.5. Green arrows indicate migration of splanchnic mesoderm cells into the pharyngeal arches. Red lines signify enriched localization of Fn1 protein at germ layer borders. Integrin α5β1 is expressed in all cell types in the pharynx at E8.5 (Chen et al., 2015) and regulates the accrual of pharyngeal mesoderm from the SHF (green arrows). **A2.** During the 10^th^ day of mouse development, integrin α5β1 and Fn1 regulate the remodeling of the plexus endothelium in the 4^th^ pharyngeal arch into the PAA. Isl1 lineages are marked in green. NC-derived cells are in blue. **B.** The expression of integrin α5 in the mesoderm regulates the differentiation of NC-derived cells into VSMCs. The deletion of integrin α5 in Mesp1 lineage, which includes the SHF, leads to defective NC-to-VSMC differentiation and results in the regression of the 4^th^ PAAs leading to IAA-B and RERSA (Liang, et al., *Dev Biol* 2014). C. Fn1 becomes upregulated in NC-derived cells adjacent to the 4th PAA ECs between E10.5 and E11.5 The expression of integrin α5β1 and Fn1 in the NC-derived cells is required for NC-to-VSMC differentiation and the stability of the 4^th^ PAA (Wang et al, 2016). The deletion of either integrin α5 or Fn1 in the NC (and the surface ectoderm) leads to IAA-B and RERSA but does not impair PAA formation. **D.** The expression integrin α5 or Fn1 in the SHF is not required for the morphogenesis of the aortic arch and its branches. **E.** The expression of integrin α5 in the endoderm and the pharyngeal endothelia is not required for the development of the aortic arch and its branches.

The differentiation of NC into VSMCs is orchestrated in part by a relay of Notch signaling transduced from the PAA endothelium to the surrounding layers of NC-derived cells ^56^. The activation of Notch signaling in the NC is required for the differentiation of NCs into VSMCs ^52, 56^. We demonstrated that this pathway was regulated by the expression of integrin α5 and fibronectin specifically by NC-derived cells at E11.5 ^31^. To test the possibility that the expression of integrin α5 in the *Isl1* lineages regulates the lateral propagation of Notch from the PAA endothelium to the adjacent NC-derived cells, we stained control and mutant sections with an antibody to Notch Intracellular Domain (NICD), an activated form of Notch. However, Notch signaling was activated comparably in controls and mutants, despite the severe deficiency in the differentiation of NC cells into VSMCs in the mutants (compare Fig. 7C2 – C4 with Fig. 7D2 – D4, arrows). These experiments indicate that the expression of integrin α5 in the pharyngeal arch mesoderm regulates the differentiation of NC cells into VSMCs independently of Notch. Furthermore, these experiments indicate that while the activation of Notch is necessary for the differentiation of NC-derived cells into VSMCs, it is not sufficient. Taken together, with our previous work ^31, 59^, studies in this manuscript demonstrate that cell-ECM interactions regulated by integrin α5β1 and Fn1 play multiple, pleiotropic, and stage-specific functions during the morphogenesis of the 4^th^ PAAs (Fig. 8).

### Combinatorial expression of integrin α5 and fibronectin from multiple lineages in the pharynx regulates the formation of the 4^th^ PAAs

The *Isl1* lineages encompass the mesoderm in the SHF and pharyngeal arches, pharyngeal endoderm, surface ectoderm, and some NC-derived cell populations, although not the NC in the pharyngeal arches ^29, 42, 59, 60^. Our previous studies indicated that the combined expression of integrin α5β1 or Fn1 in the surface ectoderm and the NC was not required for the formation of the 4^th^ PAAs ^29, 31^. However, even though PAA formation occurred normally in these mice, the 4^th^ PAAs regressed later due to defects in the differentiation of NC-derived cells into VSMCs, resulting in RERSA and IAA-B ^31^ (Fig. 8). The expression of either integrin α5β1 or Fn1 in the SHF lineage marked by the expression of the Mef2C-AHF-Cre transgene is also not required for PAA formation (Sup. Tables 1 and 2), indicating that the expression of integrin α5β1 or Fn1 in the SHF alone is not required for cardiovascular development. Consistent with these findings, the expression of integrin α5β1 in the *Mesp1* lineage or in the endothelium was not required for PAA formation ^59, 61^ (Fig. 8). Instead, the expression of integrin α5β1 in the Mesp1 lineage was required for PAA stability, and the deletion of integrin α5 in *Mesp1* lineage which includes the PAA endothelium resulted in IAA-B and RERSA (Fig. 8) ^59, 61^.

The difference in the phenotypes resulting from the deletion of integrin α5 using Mef2C-AHF-Cre and Mesp1^Cre^ are likely the result of differences in the timing of Cre expression (e.g. the later onset of Mef2C-AHF-Cre may have allowed the perdurance of integrin α5β1 protein through the stages where it’s required for mesoderm-NC interactions). Alternatively, the expression of integrin α5β1 in Mesp1 lineage-derived mesodermal cells prior to E8.5 is important for the regulation of NC cell fate in the pharyngeal arches ^59^.

Lastly, we tested whether the expression of integrin α5 in the endoderm regulated PAA formation. For these experiments, we used the constitutive Sox17^2A-iCre^ knockin mouse line, in which Cre is expressed in the endoderm and pharyngeal arch ECs (Sup. Fig. 7A – A4) ^62^. However, PAAs formed normally in α*5*^*flox/-*^; Sox17^2A-iCre^ mutants (Sup. Fig. 7B, B1, C, C1). Together, these data indicate that combinatorial expression of integrin α5 and Fn1 in the Isl1 lineages is necessary for the proper formation of the 4^th^ PAAs (Fig. 8).

## Discussion

Proper development of the 4^th^ PAAs is central to the ability of a newborn to survive and thrive (Karunamuni et al., 2014; Moon, 2008). The formation of the 4^th^ pair of the PAAs is regulated by a number of genes including *Tbx1, Pax9, Gbx2, Fgf8, Crkl, PlexinD1, and Nrp1*, e.g., ^63-67^. However, cellular mechanisms by which these genes mediate PAA formation are not well-understood. Unraveling the dynamics of EC progenitors and their descendants during PAAs formation is vital to understanding the genetic and cellular mechanisms regulating PAA formation and how they are altered in congenital heart disease.

In this manuscript, we have demonstrated that the SHF is the primary source of the PAA endothelium and that the majority of endothelial progenitors giving rise to the PAAs are already present in the SHF by E7.5. PAA progenitors exit the SHF and contribute to the PAAs over a span of about 2 days, from E7.5 to E9.5.

Lineage labeling using constitutive Cre lines Isl1^Cre^ and Mef2C-AHF-Cre led to similar labeling of the PAA endothelium, with the exception of the 3^rd^ PAA, which is labeled 50% more efficiently when Isl1^Cre^ line of mice is used. This difference likely reflects the timing of Cre expression in Isl1^Cre^ and Mef2C-AHF-Cre strains, with Mef2C-AHF-Cre lagging by about a day ^42, 48, 68^. The difference in the labeling efficiency suggests that about half of the endothelial progenitors of the 3^rd^ PAAs have exited the SHF by the time Mef2C-AHF-Cre is expressed. Our fate mapping studies show that there are differences in the timing and the extent of SHF contribution to the PAAs. In particular, if one were to use Mef2C-AHF-Cre to generate mutations, the 4^th^ PAAs could be more affected than the 3^rd^ and the 6^th^ because in the MEf2C-AHF-Cre strain, the contribution of the SHF lineage-labeled ECs to the 4^th^ PAA endothelium is the highest.

By using whole-mount imaging and quantitative analyses of EC populations in the pharyngeal arches, we previously demonstrated that the morphogenesis of the 4^th^ PAAs occurs gradually throughout the 10^th^ day of the embryonic development and entails a rapid accumulation of ECs: endothelial population in the 4^th^ pharyngeal arch increases more than three-fold in about eight hours of development, from 30 – 39 somites ^9^. This steep increase is unlikely to occur solely due to EC proliferation, and our labeling experiments show that SHF-derived cells are still being added to the 4^th^ PAA after E9.5.

Our studies show that integrin α5β1 and Fn1 are important for initial recruitment of SHF-derived ECs into the 4^th^ pharyngeal arches, and that in the absence of integrin α5 or Fn1 in the Isl1 lineages results in EC deficiency up to 32 – 34 somite stage. Despite the initial EC deficiency in the 4^th^ arch, EC numbers recover in integrin α5^f/-^; Isl1^Cre/+^ and Fn1^f/-^; Isl1^Cre/+^ mutants by the 34 – 35 somite stage. We demonstrate that the recovery of EC cell numbers in the pharyngeal arches is not due to compensation from an alternative endothelial source. Instead, we show that the proliferation index of plexus endothelium is 2-fold higher than that of ECs in the 4^th^ PAA (Fig. 3O). This difference in the proliferation index is maintained in the mutants (Fig. 3O). We hypothesize that since the majority of ECs is in the plexus at 32 – 33 somites in the mutants, their proliferative advantage over ECs in the PAA allows the EC number in the mutant arches to recover by the end of E10.5.

In spite of the recovery of EC populations in the 4^th^ arches, the 4^th^ PAAs were either thin or absent in 50% of all the 4^th^ arches by 36 – 39 somite stages ^29^. Our regression analysis showed that the rearrangement of the 4^th^ pharyngeal arch ECs into the PAA was not dependent on the number or density of ECs in the 4^th^ arch. These data suggest that the remodeling of the uniform endothelial plexus into the PAA in the 4^th^ arch is mediated by factors extrinsic to the endothelium. The disruption of the remodeling in our mutants indicates an essential role for cell – ECM interactions in this process.

The *Isl1* lineage encompasses multiple cell types within the pharynx including pharyngeal epithelia, mesoderm, and a population of NC cells in the cardiac outflow tract ^60, 68^. Pharyngeal endoderm and the ectoderm are important signaling centers regulating intercellular communications among the germ layers composing the arches during the morphogenesis of cardiopharyngeal organs and structures ^27, 69, 70^. Modulation of the extracellular microenvironment within the pharynx is essential for the development cardiovascular system ^29, 31, 59, 63, 64, 71-81^. The expression of *Fn1*, an essential ECM glycoprotein, is highly enriched in the pharyngeal epithelia ^28, 29^, and our studies show that signaling by Fn1 in the Isl1 lineages is important for the accrual of SHF-derived cells to the pharyngeal mesenchyme, and for the formation of the 4^th^ PAAs. In the latter step, signaling by Fn1 in tissues extrinsic to the pharyngeal arch endothelium regulates the remodeling of endothelial plexus into the PAA in the 4^th^ arch. While Fn1 expression in the NC regulates PAA stability after the 4^th^ PAA has formed ^29, 31^ (Fig. 8).

In this manuscript, we investigated the tissues wherein signaling by Fn1 is important for PAA formation. Integrin α5β1 is a major Fn1 receptor during embryogenesis ^28-30, 34, 38, 39, 41^, and the deletion of integrin α5 or Fn1 in the Isl1 lineages results in identical phenotypes ^29^. To determine the cell type(s) in which signaling by Fn1 regulates PAA formation, we ablated integrin α5 in each of the tissues comprising Isl1 lineage individually or in combination. The deletion of integrin α5 in the SHF (Mef2C-AHF-Cre strain), the entire anterior mesoderm (Mesp1^Cre^), the NC (Wnt1-Cre2, P3Pro-Cre), the NC and surface ectoderm (TFAP2α^IresCre^), or the endoderm and endothelia (Sox17^2A-iCre^) resulted in normal PAA formation (this study and ^29, 31, 59^). Therefore, we conclude that combinatorial signaling by integrin α5β1 from pharyngeal endoderm, mesoderm, and the surface ectoderm is essential to mediate the formation of the 4^th^ PAAs. While signaling in the mesoderm and the neural crest is important for PAA stability (Fig. 8).

The PAAs form within the neural crest-derived pharyngeal mesenchyme and the PAA endothelium induces the differentiation of the adjacent NC-derived cells into VSMCs ^56^. Despite the initial delay in the formation of the 4^th^ PAAs, the size of PAAs in integrin α5^f/-^; Isl1^Cre/+^ mutants recovers by E11.5. At this time, we observed a profound deficiency in the expression of αSMA by NC-derived cells around the PAAs in the mutants. Deficiency in VSMC differentiation causes vessel regression ^31, 52, 57, 82, 83^, consistent with our finding that integrin α5^f/-^; Isl1^Cre/+^ and Fn1^f/-^; Isl1^Cre/+^ mutants develop IAA-B and RERSA; defects that are caused by the aberrant morphogenesis of the left and right 4^th^ PAAs, respectively ^29^. Our previous studies using integrin α5^f/-^; Mesp1^Cre/+^ mice demonstrated that the expression of integrin α5 in the mesoderm regulates NC differentiation into VSMCs without affecting PAA formation, and integrin α5^f/-^; Mesp1^Cre/+^ mice develop IAA-B and RERSA ^59^ (Fig. 8). Thus, the roles of integrin α5β1 and Fn1 in the formation of the 4^th^ PAAs are separate from their roles in the differentiation of NC-derived cells into VSCMs ^31^.

Mechanisms that lead to IAA-B are complex but generally arise due to either of the following two broad categories of defects: a) defects in the formation of the left 4^th^ PAA or b) defects in the stability of an otherwise well-formed 4^th^ PAA. NC ablation studies in chick and genetic manipulation of the neural crest demonstrate that NC is not required for PAA formation ^51, 52^. Even in the extreme case of neural crest ablation, PAAs form ^51^. However, defective differentiation of NC-derived cells into VSMCs leads to PAA regression resulting in various malformations in the aortic arch and its branches, including IAA-B and RERSA ^31, 52, 82^. Our studies show that the expression of integrin α5β1 in the pharyngeal mesoderm and the NC are required for NC-to-VSMC differentiation, and the expression of integrin α5β1 in either of these lineages alone is not sufficient for this process ^31, 59^.

Defects in the formation of the 4^th^ PAAs often occur in conjunction with 22q11 deletion syndromes ^84^. Cumulatively, four prospective studies found that between 40 and 90% of interrupted aortic arch type B (IAA-B) cases diagnosed in fetuses, neonates, and children can be attributed to deletions in the 22q11 region ^85^. Studies using *Tbx1*^*+/-*^ mice that model 22q11 deletion syndrome indicated that defective formation of the left 4^th^ PAA underlies IAA-B in these patients ^65, 86, 87^. Intriguingly, several independent publications demonstrated that *Tbx1* regulates the expression of integrins and extracellular matrix (ECM) components, and showed that defects in cell-ECM interactions downstream of *Tbx1* precede pathological sequelae and cardiovascular defects in *Tbx1* mutants ^71-73^. Interestingly, about 50% of Tbx1^+/-^ mice recover from the initial defect in PAA formation, and are viable and fertile ^50^; However, this recovery can be impeded by the reduction in the expression of Fn1 ^49^. Thus, alterations in cell-ECM interactions and pharyngeal ECM microenvironment may underlie lethal AAA defects in 22q11 deletion syndrome downstream of *Tbx1*.

The significance of our work lies in the identification of cellular dynamics regulating PAA formation and the intricate temporal and cell-type specific roles of cell-ECM interactions in the regulation of aortic arch morphogenesis at multiple steps of its formation and remodeling (Fig. 8).

## Supporting information

ExtendedMethods

## Nonstandard Abbreviations and Acronyms in the Alphabetical Order

AAA: aortic arch artery
CHD: congenital heart disease
ECs: endothelial cells
Fn1: fibronectin
IAA-B: interrupted aortic arch type B
Itga5: integrin α5
PAA: pharyngeal arch arteries
RERSA: retro-esophageal right subclavian artery
SHF: second heart field
VEGFR2: vascular endothelial growth factor receptor 2

## Acknowledgements

We thank Richard Hynes, Chenleng Cai, Sylvia Evans, Benoit Bruneau, Heicko Lickert, and Hongkui Zeng for the gift of mouse strains, and Brianna Alexander and Nathan Astrof for careful reading of the manuscript.

## Supplemental Materials

Expanded Materials & Methods

Online Figures 1 – 7

---

**Supplemental Figure 1. Delayed formation of the 4**^**th**^ **PAAs in mutants lacking integrin α5 or Fn1 in the Isl1 lineages. A – B.** Quantification of total endothelial cell numbers in the 4^th^ arches of controls and mutants show similar phenotypes among integrin α5^flox/-^; Isl1^Cre/+^ and Fn1^flox/-^; Isl1^Cre/+^ embryos. Note the recovery of endothelial cell numbers at 36 – 40-somite stage. **C.** Combined data comparing endothelial populations of controls and integrin α5^flox/-^; Isl1^Cre/+^ and Fn1^flox/-^; Isl1^Cre/+^ mutant embryos at 33 – 34-somite stages. **D.** The proportion of pharyngeal ECs in the plexus is increased in the mutants relative to controls. **E.** The proportion of endothelial cells in the PAA is decreased in the mutants relative to controls. Note, 7 of 16 mutants did not have PAAs (0% endothelial cells in the 4^th^ PAA).

**Supplemental Figure 2. The prevalence of cell death, as assayed by the presence of cleaved caspase 3 or TUNEL signals, was similar in controls and mutants. A, C.** Controls. **B, D.** Mutants. All scale bars are 100 μm.

**Supplemental Figure 3. The differentiation of SHF-derived cells into endothelial cells is not affected by the deletion of integrin α5 in the Isl1 lineages.** Whole-mount staining, confocal imaging and 3D reconstruction through the pharyngeal regions of control (**A, C**) and mutant (**B, D**) embryos. **A – B.** Sagittal optical sections through E9.5 embryos: The majority of VEGFR2^+^ cells express GFP and ERG in the 3^rd^ PAA and in a more posterior mesenchyme (arrows) in controls and mutants. **C – D.** E10.5 embryos. 3D reconstruction through the pharyngeal region. Open chevrons mark the 4^th^ PAAs. Note the presence of a very thin PAA in the mutant (**D-D1**). All VERGFR2^+^ cells are Pecam1^+^ in control and in the mutant with defective 4th PAA. Scale bars are 50 μm.

**Supplemental Figure 4. Quantification of SHF-derived cells in pharyngeal mesenchyme and splanchnic mesoderm.** E9.5 Isl1^Cre/+^; Rosa^nTnG/+^ embryos were stained with anti-GFP antibodies, cleared in BABB, and imaged through the entire pharyngeal region using 25x silicone oil objective N.A. 1.05 and Nikon confocal microscope. 3D reconstruction and surfacing were done using IMARIS. **A.** Pharyngeal mesenchyme in the 1^st^ and 2^nd^ pharyngeal arches was surfaced. Dashed line marks the plane of transverse optical section shown in **A1.** GFP^+^ cells within the pharyngeal mesenchyme (yellow) were quantified using the spot function in IMARIS in the entire volume marked by the yellow surfaces in **A. B –B4.** Splanchnic mesoderm within the dorsal pericardial wall was surfaced in pink, and pharyngeal mesenchyme was surfaced in yellow. **B.** Ventral view. **B1.** Right-side view. Dashed line indicates the plane of section shown in **B3. B2.** A slanted, sagittal/coronal view to visualize both the splanchnic mesoderm and pharyngeal mesenchyme. **B3.** GFP^+^ cells in the splanchnic mesoderm (pink) and in the posterior pharyngeal mesenchyme (yellow) were quantified using the spot function in IMARIS throughout the entire volume shown in **B.**

**Supplemental Figure 5. Step-wise changes in the configuration of the 4**^**th**^ **arch endothelium on the 10**^**th**^ **day of mouse embryonic development.** Control embryos were stained using antibody to Pecam1. Endothelial cells in the 4^th^ pharyngeal arch were surface-rendered yellow using IMARIS. First row, 30-somite stage. Endothelium in the 4^th^ arch is in the form of a plexus of small blood vessels. Second row 33-somite embryo. A small PAA is seen forming. Red starts mark spaces amidst the interconnected plexus vessels and the small PAA. Third row, 33-somite stage. A large PAA is seen with connecting plexus vessels. Spaces (marked by the red stars) are still seen. Fourth row – 36 – 40 – somite stage. A large PAA is present in the 4^th^ arch by the evening of the 10^th^ day. DA – dorsal aorta. PAAs are numbers. All scale bars are 50 μm.

**Supplemental Figure 6. The expression of integrin α5β1 in the Isl1 lineages regulates the differentiation of neural crest-derived cells into VSCMs at E11.5** Activation of Notch in the pharyngeal arch mesenchyme is not altered in the mutants. Coronal section through the pharyngeal regions of Control (**A**) and Mutant (**B**) embryos were stained to detect green fluorescent protein (GFP, green) marks the Isl1 lineages; Notch intracellular domain (NICD, orange) is used as the readout of active Notch signaling; and a smooth muscle actin (αSMA, blue) marks smooth muscle cells. PAAs are numbered. The magnification is the same in all panels.

**Supplemental Figure 7. PAA formation is not affected when integrin α5 is ablated using Sox17**^**2A-iCre**^ **knock-in strain.** Whole-mount pictures were taken following India Ink injections into the hearts of controls and mutants isolated at E10.5. Magnification is the same in all panels.

