## ExtendedMethods for "Cell – ECM interactions play distinct and essential roles at multiple stages during the development of the aortic arch"

**Animals** C57BL/6J mice (cat # 0664) were purchased from Jackson Laboratories. Integrin $\alpha 5^{+/-}$ and integrin $\alpha 5^{flox/flox}$ mice were gifts from Dr. Richard Hynes (van der Flier et al., 2010; Yang and Hynes, 1996). Fn1$^{flox/flox}$ mice were a gift from Reinhardt Fassler (Sakai et al., 2001). *Isl1*$^{Cre}$ and *Isl1*$^{MerCreMer}$ knock-in mice were gifts from Dr. Sylvia Evans (Cai et al., 2003; Sun et al., 2007), Mef2C-AHF-DreERT2 transgenic mice were a gift from Dr. Benoit Bruneau (Devine et al., 2014), Sox17$^{2A\text{-}iCre}$ knock-in mice were a gift from Heicko Lickert (Engert et al., 2009), Rosa$^{mTmG}$ mice, Gt (ROSA)26Sortm4(ACTB-tdTomato,–EGFP) (Muzumdar et al., 2007) and B6;129S6-*Gt(ROSA)26Sor*$^{tm1(CAG\text{-}tdTomato*,\text{-}EGFP*)Ees}$/J, known as ROSA$^{nT\text{-}nG}$ mice, generated by Justin Prigge and Ed Schmidt at Montana State University, were purchased from Jackson labs; in this strain, two distinct nuclear localization signals derived from human SRm160 protein (Wagner et al., 2003) were fused with tdTomato and EGFP sequences.  Dre reporter mice were a gift from Dr. Hongkui Zeng (Madisen et al., 2015). All Cre-expressing strains were crossed into C57BL6/J background for at least 10 generations. Integrin $\alpha 5^{flox/flox}$ were and Fn1$^{flox/flox}$ mice were originally on a mixed 129/C57BL6/J genetic background. These strains were crossed into ROSA reporter strains (each of which was crossed into C57BL6/J background by the Jackson Labs) and maintained in our lab for the past 10 years.

Mice and embryos were genotyped according to published protocols. Embryos were staged by counting somites. Mice were housed in an AAALAC-approved barrier facility. All experimental procedures were approved by the Institutional Animal Care and Use Committee of Rutgers University and conducted in accordance with the Federal guidelines for the humane care of animals.

**Whole Mount Immunofluorescence staining** Labeling with BrdU, TUNEL, and staining with antibodies against cleaved caspase 3 were performed as described in (Chen et al., 2015). Embryos were isolated at specified days of development, fixed in 4% paraformaldehyde (PFA) at $4^oC$ overnight, rinsed in PBS and either used immediately, or dehydrated through a series of 25%, 50%, and 75% of methanol dilutions in PBS, then washed three times with 100% methanol and stored in 100% methanol at $-20^oC$ until use. The following antibodies were used $1^o$ antibodies: anti-Pecam1 (1:200, BD Pharmingen, cat #550274), anti-VEGFR2 (1:200, R&D, cat #AF644), anti-ERG (1:1000, Abcam, cat# ab214341), anti-GFP (1:300, Aves, cat #GFP1020). All $2^o$ antibodies were from Invitrogen and used at the dilution of 1:300. Nuclei were stained using DAPI (1:1000 dilution of 5 mg/ml stock diluted in $H_2O$, Sigma, cat #32670-5MG-F). Following washes, stained embryos were embedded into 1% agarose (Bio-Rad Laboratories, cat #1613101), cleared using Benzyl Alcohol (Sigma, cat # B-1042)/ Benzyl Benzoate (Sigma, cat #B-6630) and placed between two #1.5 coverslips (VWR, cat #16004-312) separated by a rubber spacer (Grace Bio Labs, cat # 664113).

**Imaging and quantifications** Confocal imaging was done using Nikon A1R microscopes with 20x CFI Apo LWD Lambda S water immersion objective (MRD77200) or 25x CFI Plan Apo Lambda S silicone oil objectives (MRD73250). 3D reconstructions, surfacing, and quantifications were performed using IMARIS software (Bitplane, USA) (Ramirez and Astrof, 2020; Wang et al., 2017). The total number of ECs was determined by quantification of $ERG^+VEGFR2^+$ ECs. The number of ECs in PAAs was determined by masking the PAA VEGFR2$^{+}$ surface and quantifying the number of ERG$^{+}$ nuclei within. Detailed, step-by-step imaging and quantification procedures are described in (Ramirez and Astrof, 2020). For quantification of SHF-derived cells in pharyngeal arches and pharyngeal endothelium, nuclei of SHF-derived cells were marked by the expression of GFP through the use of ROSA$^{nT\text{-}nG}$ reporter mice. The precise areas analyzed in E9.5 embryos are shown in Sup. Fig. 4. For the quantification of SHF-derived cells at E10.5, embryos were stained to detect VEGFR2, ERG, nuclear GFP, and DAPI, and surfacing and quantification were carried out as described in (Ramirez and Astrof, 2020).

**References associated with the Methods**

Cai, C.L., Liang, X., Shi, Y., Chu, P.H., Pfaff, S.L., Chen, J., Evans, S., 2003. Isl1 identifies a cardiac progenitor population that proliferates prior to differentiation and contributes a majority of cells to the heart. Dev Cell 5, 877-889.

Chen, D., Wang, X., Liang, D., Gordon, J., Mittal, A., Manley, N., Degenhardt, K., Astrof, S., 2015. Fibronectin signals through integrin alpha5beta1 to regulate cardiovascular development in a cell type-specific manner. Dev Biol 407, 195-210.

Devine, W.P., Wythe, J.D., George, M., Koshiba-Takeuchi, K., Bruneau, B.G., 2014. Early patterning and specification of cardiac progenitors in gastrulating mesoderm. eLife 3.

Engert, S., Liao, W.P., Burtscher, I., Lickert, H., 2009. Sox17-2A-iCre: a knock-in mouse line expressing Cre recombinase in endoderm and vascular endothelial cells. Genesis 47, 603-610.

Madisen, L., Garner, A.R., Shimaoka, D., Chuong, A.S., Klapoetke, N.C., Li, L., van der Bourg, A., Niino, Y., Egolf, L., Monetti, C., Gu, H., Mills, M., Cheng, A., Tasic, B., Nguyen, T.N., Sunkin, S.M., Benucci, A., Nagy, A., Miyawaki, A., Helmchen, F., Empson, R.M., Knopfel, T., Boyden, E.S., Reid, R.C., Carandini, M., Zeng, H., 2015. Transgenic mice for intersectional targeting of neural sensors and effectors with high specificity and performance. Neuron 85, 942-958.

Muzumdar, M.D., Tasic, B., Miyamichi, K., Li, L., Luo, L., 2007. A global double-fluorescent Cre reporter mouse. Genesis 45, 593-605.

Ramirez, A., Astrof, S., 2020. Visualization and Analysis of Pharyngeal Arch Arteries using Whole-mount Immunohistochemistry and 3D Reconstruction. J. Vis. Exp. 157, e60797.

Sakai, T., Johnson, K.J., Murozono, M., Sakai, K., Magnuson, M.A., Wieloch, T., Cronberg, T., Isshiki, A., Erickson, H.P., Fassler, R., 2001. Plasma fibronectin supports neuronal survival and reduces brain injury following transient focal cerebral ischemia but is not essential for skin-wound healing and hemostasis. Nat Med 7, 324-330.

Sun, Y., Liang, X., Najafi, N., Cass, M., Lin, L., Cai, C.L., Chen, J., Evans, S.M., 2007. Islet 1 is expressed in distinct cardiovascular lineages, including pacemaker and coronary vascular cells. Dev Biol 304, 286-296.

van der Flier, A., Badu-Nkansah, K., Whittaker, C.A., Crowley, D., Bronson, R.T., Lacy-Hulbert, A., Hynes, R.O., 2010. Endothelial alpha5 and alphav integrins cooperate in remodeling of the vasculature during development. Development 137, 2439-2449.

Wagner, S., Chiosea, S., Nickerson, J.A., 2003. The spatial targeting and nuclear matrix binding domains of SRm160. Proc Natl Acad Sci U S A 100, 3269-3274.

Wang, X., Chen, D., Chen, K., Jubran, A., Ramirez, A., Astrof, S., 2017. Endothelium in the pharyngeal arches 3, 4 and 6 is derived from the second heart field. Dev Biol 421, 108-117.

Yang, J.T., Hynes, R.O., 1996. Fibronectin receptor functions in embryonic cells deficient in alpha 5 beta 1 integrin can be replaced by alpha V integrins. Mol Biol Cell 7, 1737-1748.
