## Supplemental Figure 1 for "Cell – ECM interactions play distinct and essential roles at multiple stages during the development of the aortic arch"

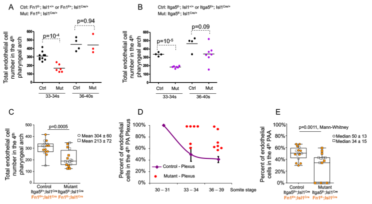


**Supplemental Figure 1.** Delayed formation of the 4th PAAs in mutants lacking integrin $\alpha5$ or Fn1 in the Isl1 lineages. **A – B.** Quantification of total endothelial cell numbers in the 4th arches of controls and mutants show similar phenotypes among integrin $\alpha5^{flox/-}$; $Isl1^{Cre/+}$ and $Fn1^{flox/-}$; $Isl1^{Cre/+}$ embryos. Note the recovery of endothelial cell numbers at 36 – 40-somite stage. **C.** Combined data comparing endothelial populations of controls and integrin $a5^{flox/-}$; $Isl1^{Cre/+}$ and $Fn1^{flox/-}$; $Isl1^{Cre/+}$ mutant embryos at 33 – 34-somite stages. **D.** The proportion of pharyngeal ECs in the plexus is increased in the mutants relative to controls. **E.** The proportion of endothelial cells in the PAA is decreased in the mutants relative to controls. Note, 7 of 16 mutants did not have PAAs (0% endothelial cells in the 4th PAA).
