## Supplemental Figure 2 for "Cell – ECM interactions play distinct and essential roles at multiple stages during the development of the aortic arch"

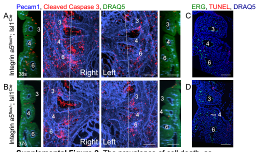


**Supplemental Figure 2.** The prevalence of cell death, as assayed by the presence of cleaved caspase 3 or TUNEL signals, was similar in controls and mutants. **A, C.** Controls. **B, D.** Mutants. All scale bars are 100 μm.
