## Supplemental Figure 3 for "Cell – ECM interactions play distinct and essential roles at multiple stages during the development of the aortic arch"

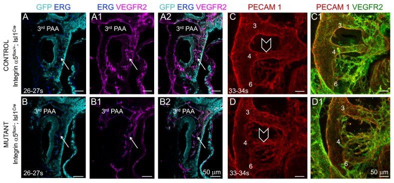


**Supplemental Figure 3**. **The differentiation of SHF-derived cells into endothelial cells is not affected by the deletion of integrin α5 in the Isl1 lineages.** Whole-mount staining, confocal imaging and 3D reconstruction through the pharyngeal regions of control (**A, C**) and mutant (**B, D**) embryos. **A – B.** Sagittal optical sections through E9.5 embryos: The majority of $VEGFR2^+$ cells express GFP and ERG in the 3rd PAA and in a more posterior mesenchyme (arrows) in controls and mutants. **C – D.** E10.5 embryos. 3D reconstruction through the pharyngeal region. Open chevrons mark the 4th PAAs. Note the presence of a very thin PAA in the mutant (**D-D1**). All $VERGFR2^+$ cells are $Pecam1^+$ in control and in the mutant with defective 4th PAA. Scale bars are 50 μm.
