## Supplemental Figure 4 for "Cell – ECM interactions play distinct and essential roles at multiple stages during the development of the aortic arch"

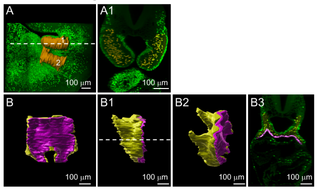


**Supplemental Figure 4.** Quantification of SHF-derived cells in pharyngeal mesenchyme and splanchnic mesoderm. E9.5 $Isl1^{Cre/+}$; $Rosa^{nTnG/+}$ embryos were stained with anti-GFP antibodies, cleared in BABB, and imaged through the entire pharyngeal region using 25x silicone oil objective N.A. 1.05 and Nikon confocal microscope. 3D reconstruction and surfacing were done using IMARIS. **A.** Pharyngeal mesenchyme in the $1^{st}$ and $2^{nd}$ pharyngeal arches was surfaced. Dashed line marks the plane of transverse optical section shown in **A1**. $GFP^+$ cells within the pharyngeal mesenchyme (yellow) were quantified using the spot function in IMARIS in the entire volume marked by the yellow surfaces in **A. B –B4**. Splanchnic mesoderm within the dorsal pericardial wall was surfaced in pink, and pharyngeal mesenchyme was surfaced in yellow. **B.** Ventral view. **B1.** Right-side view. Dashed line indicates the plane of section shown in **B3. B2.** A slanted, sagittal/coronal view to visualize both the splanchnic mesoderm and pharyngeal mesenchyme. **B3**. $GFP^+$ cells in the splanchnic mesoderm (pink) and in the posterior pharyngeal mesenchyme (yellow) were quantified using the spot function in IMARIS throughout the entire volume shown in **B**.
