## Supplemental Figure 5 for "Cell – ECM interactions play distinct and essential roles at multiple stages during the development of the aortic arch"

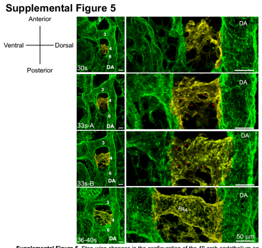


**Supplemental Figure 5**. Step-wise changes in the configuration of the 4th arch endothelium on the 10th day of mouse embryonic development. Control embryos were stained using antibody to Pecam1. Endothelial cells in the 4th pharyngeal arch were surface-rendered yellow using IMARIS. First row, 30-somite stage. Endothelium in the 4th arch is in the form of a plexus of small blood vessels. Second row 33-somite embryo. A small PAA is seen forming. Red starts mark spaces amidst the interconnected plexus vessels and the small PAA. Third row, 33-somite stage. A large PAA is seen with connecting plexus vessels. Spaces (marked by the red stars) are still seen. Fourth row – 36 – 40 – somite stage. A large PAA is present in the 4th arch by the evening of the 10th day. DA – dorsal aorta. PAAs are numbers. All scale bars are 50 μm.
