## Supplemental Figure 6 for "Cell – ECM interactions play distinct and essential roles at multiple stages during the development of the aortic arch"

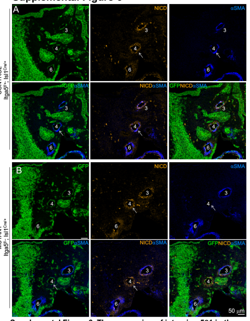


**Supplemental Figure 6. The expression of integrin α5β1 in the Isl1 lineages regulates the differentiation of neural crest-derived cells into VSCMs at E11.5** Activation of Notch in the pharyngeal arch mesenchyme is not altered in the mutants. Coronal section through the pharyngeal regions of Control (**A**) and Mutant (**B**) embryos were stained to detect green fluorescent protein (GFP, green) marks the Isl1 lineages; Notch intracellular domain (NICD, orange) is used as the readout of active Notch signaling; and α smooth muscle actin (αSMA, blue) marks smooth muscle cells. PAAs are numbered. The magnification is the same in all panels.
