## Supplemental Figure 7 for "Cell – ECM interactions play distinct and essential roles at multiple stages during the development of the aortic arch"

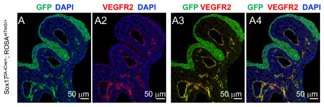


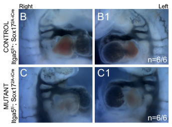


**Supplemental Figure 7. PAA formation is not affected when integrin α5 is ablated using Sox17$^{2A\text{-}iCre}$ knock-in strain. A.** Coronal sections through the pharyngeal region of E10.5 embryos showing the fate map of Sox17 lineage in Sox17$^{2A\text{-}iCre}$ mice. **B – C.** Whole-mount pictures were taken following India Ink injections into the hearts of control (**B – B**1) and mutant (**C – C 1**) embryos isolated at E10.5. Magnification is the same in all panels.
