## Supplementary material for "Cell – ECM interactions play distinct and essential roles at multiple stages during the development of the aortic arch": Tables

**Supplemental Table 1.** Itga5$^{f/f}$ x Itga5$^{+/-}$; Mef2C-AHF-Cre

| Genotype | # Observed at Weaning |
|---|---|
| Itga5$^{f/-}$ ; Mef2C-AHF-Cre+ | 14 (22%) |
| Itga5 $^{f/-}$ | 12 (19%) |
| Itga5$^{f/+}$ ; Mef2C-AHF-Cre+ | 18 (29%) |
| Itga5 $^{f/+}$ | 19 (30%) |
| Total | 63 |

**Supplemental Table 2**. Fn1$^{f/f}$ x Fn1$^{+/}$-; Mef2C-AHF-Cre

| Genotype | # Observed at Weaning |
|---|---|
| Fn1$^{f/-}$ ; Mef2C-AHF-Cre+ | 23 (31%) |
| Fn1 $^{f/-}$ | 15 (20%) |
| Fn1$^{f/+}$ ; Mef2C-AHF-Cre+ | 17 (23%) |
| Fn1 $^{f/+}$ | 19 (26%) |
| Total | 74 |
